# Coordination of focal adhesion nanostructure and mechano-signaling drives cardiomyocyte differentiation

**DOI:** 10.1101/2023.11.13.566796

**Authors:** Jingwei Xiao, Xueying Zhong, Jing Wen Ang, Darren Chen Pei Wong, Chang Jie Mick Lee, Roger S-Y Foo, Pakorn Kanchanawong, Boon Chuan Low

**Affiliations:** Mechanobiology Institute Singapore, National University of Singapore, Singapore; Department of Biological Sciences, National University of Singapore, Singapore; Department of Medicine, Yong Loo Lin School of Medicine, National University of Singapore, Singapore; Cardiovascular Research Institute, National University Healthcare Systems, Singapore; Department of Biomedical Engineering, National University of Singapore, Singapore; NUS College, National University of Singapore, Singapore

## Abstract

Focal adhesion (FA) organization and signaling are essential for cell growth and differentiation. However, the molecular mechanism that coordinates the FA signaling with cardiomyocyte differentiation has not been fully understood. Here, we provide empirical evidence that BNIP-2, a BCH-domain-containing protein, is the organizer of FA nanostructure that potentiates FA signaling and cell traction force transmission. Mechanistically, BNIP-2 serves as a scaffold for focal adhesion kinase (FAK), paxillin and vinculin to control their molecular organization and assembly/disassembly within FAs. Constitutively active phosphomimetic mutant FAK Y397D shows enhanced binding to BNIP-2, whereas the depletion of BNIP-2 reduces FAK phosphorylation and interaction between FAK and paxillin. Using H9c2 myoblasts and human embryonic stem cells as model systems, we show that BNIP-2 depletion results in aberrant FA dynamics with impairment of traction force, and changes in signature target genes, hereby impeding cardiomyocyte differentiation. BNIP-2 regulation of FA organization and dynamic is therefore pivotal to the mechanotransduction in cardiomyocyte differentiation, shedding new light to how FA-transduced force modulates cell growth and differentiation.

## Introduction

Cells experience various forces during embryo development, such as stretching forces and pressure from the surroundings^1^. These forces influences the molecular processes underlying various cellular processes, such as cell migration, proliferation, and differentiation^2, 3^. The regulation of mechanical forces on mediating heart development has been elucidated in several animal models, such as mice, chicken, and zebrafish^4^. Of which, integrin-mediated adhesions are the key regulators of force transmission^5^. The perturbation of integrin-mediated force transmission is strongly associated with cardiovascular pathological remodelings, such as fibrosis and hypertrophy ^6–8^. Furthermore, the cardiovascular diseases is often noted with increased myocardial stiffness, suggesting that the regulation of mechanical cues from the tissue microenvironment is critical for heart development and pathologies^9^. Despite our increasing understanding of these links, the detailed molecular mechanisms underlying mechanosensing and mechanotransduction in heart tissues remain to be further investigated.

While, the linkage between heart contractility and cardiac development is first demonstrated in the study of myosin light chain 2a knockout in mouse embryos^10, 11^, the control of cell differentiation by extracellular mechanical signals is dependent on the force transmission through focal adhesion (FA) signaling. FAs are integrin-based cell adhesion structures that connect the actin cytoskeleton to extracellular matrix (ECM) via integrins and associated proteins, thus playing essential roles in transducing extracellular mechanical signals into intracellular signaling^12, 13^. This plasma membrane-associated multi-protein complexes comprise three major nanoscale compartments: the integrin signaling layer (ISL), the force-transduction layer, and the actin regulatory layer^12^. Such integrated multi-layered structure is thought to form the structural basis of the ‘molecular clutch’ function of FAs to transduce traction forces from extracellular microenvironment into the intracellular sarcomeres, which are the key regulator of muscle cell contractility and maintain the force homeostasis in cardiomyocytes ^14–18^. Focal adhesion kinase (FAK) is a key signaling component of FAs that interact directly and indirectly with integrins and various adapters such as paxillin and vinculin^19, 20^. FAK governs cardiomyocyte growth in size, and cardiac hypertrophy occurs in mice with FAK inactivation^21–23^.

Actin cytoskeleton remodeling mediated by force generation and transmission within the cell is regulated by Rho-family small GTPases and its potent downstream effectors ROCK and mDia^24^. The tension generated by actomyosin contractility is known to mediate the molecular kinetics of FA proteins. The inhibition of myosin II contractility results in the increased dissociation of vinculin and slower dissociation of paxillin and zyxin from FAs^25^. In the meantime, tension promotes force transmission by stabilizing active vinculin conformation^26^. High tension across vinculin is associated with adhesion assembly and growth, whereas low tension is accompanied by FA disassembly or rear FA sliding^27^. The regulation of FA dynamics and signaling is crucial for cellular force transmission, and these pathways are known to contribute to direct heart development and cardiac pathologies^9, 28^.

BCL2/adenovirus E1B 19 kDa protein-interacting protein 2 (BNIP-2) is a versatile signaling scaffold expressed in most tissues, with high expression observed in smooth muscle, heart muscle, and skeletal muscle^29^. BNIP-2 is essential for regulating cellular behaviors (e.g. morphogenesis, motility) through its interactions with multiple proteins, and is also responsible for regulating skeletal muscle cell differentiation and Rho-dependent force transmission in breast cancer cells^30–33^. However, the detailed mechanism by which BNIP-2 regulates cardiomyocyte differentiation is unknown. We previously showed that BNIP-2 reduction impedes cardiomyogenesis^29^. Therefore, we hypothesize that BNIP-2 plays a role in regulating mechanotransduction to direct cardiomyocyte differentiation.

In this study, we delineated the regulatory role of BNIP-2 on mechanotransduction in FAs by maintaining the integration of FA structure and interactions among FA proteins to coordinate their activities. BNIP-2 functions in differentiating human embryonic stem cells and H9c2 myoblasts into cardiomyocytes. Depletion of BNIP-2 results in the loss of FAK/paxillin organization and signaling accompanied by the changes in FA dynamics and impaired traction forces, henceforth impeding cardiomyocyte differentiation. Our results collectively suggest that BNIP-2 contributes to the regulation of FA nanostructure and FA mechano-signaling to direct cell differentiation. Our findings are key to understanding mechanotransduction in cardiac development, and could shed lights to tissue growth in other organ development.

## Results

### BNIP-2 knockdown reduces mechanotransduction and impedes H9c2 cardiac differentiation

Stiffness from the ECM is a crucial mechanical cue supporting and directing heart development in embryos^34^. Traction forces exerted by cardiomyocytes onto ECM can be sensed by neighbouring tissues to promote heart development^11, 35^. Past studies have shown that BNIP-2 promotes skeletal muscle cell differentiation and regulates Rho-dependent force transmission in breast cancer cells^30, 31, 33^. Recently, we showed that BNIP-2 depletion inhibits cardiogenesis^29^. As FAs are key structures that mediate force transmission between cells and the surrounding environment^12, 17, 36^, we hypothesize that BNIP-2 modulates mechanotransduction through FAs to direct cell differentiation in cardiac myoblasts. Previous studies have shown that depletion of focal adhesion kinase (FAK) results in cardiac defects^23^ and the dysfunction of the cardiovascular system in mice (Mouse Genome Database, **Figure S1**a, Supporting Information)^37, 38^. Using ctnt and myl2v as molecular markers for cardiomyocyte differentiation, we examined the effect of FAK inhibition on H9c2 cells, an immortalized myoblast cell line derived from the left ventricle of embryonic rats ^39, 40^. Inhibition of FAK impeded the differentiation of H9c2 myoblasts. Similarly, BNIP-2 knockdown phenocopied the effect of FAK inhibition on H9c2 cardiac differentiation (**Figure 1**a and Figure S1b, Supporting Information).

**Figure 1.**
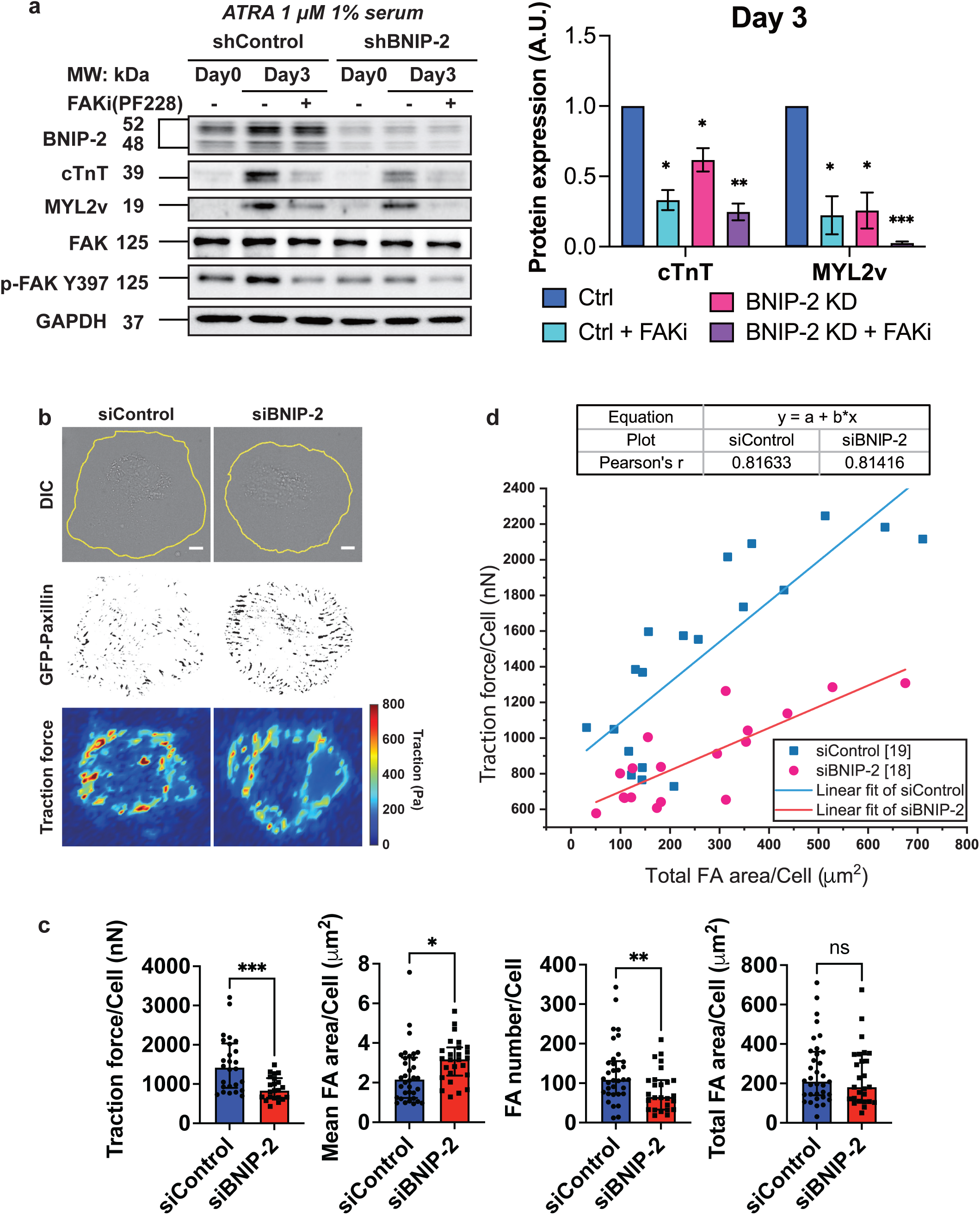
BNIP-2 knockdown reduces mechanotransduction and impedes H9c2 cardiac differentiation. **a**, Immunoblot analyses of BNIP-2, cardiac troponin-T (cTnT), ventricular myosin light chain-2 (MYL2v), FAK, phosphorylated FAK Y397 in undifferentiated (Day 0) or differentiated (Day 3) under the control or BNIP-2 knockdown H9c2 cells (left panel). Where indicated, cells were treated with 5 μM FAK inhibitor and 1 μM ATRA in low serum medium (1% FBS) for three days. BNIP-2 presents itself as different isoforms likely due to post transcriptional or/and post-translational modifications. Level of cTnT and MYL2v relative to Day 3 cells under control condition (right graph). *n*=3 independent experiments. **b**, Images showing traction force measurements of H9c2 cells expressing both siRNA targeting BNIP-2 and GFP-paxillin. The deformation is calculated as displacements of fluorescent beads coated on PDMS gel. Scale bar, 10 μm. **c**, Quantification of traction forces and morphometric analyses of FAs. The total traction forces, mean area of FAs, FA numbers, and total FA area of siBNIP-2 H9c2 myoblasts are shown (left to right). Each data point represents independent cells examined over four independent experiments. Traction force measurements, *n* = 26 (siControl) and 22 (siBNIP-2) cells; Morphometric analysis of FAs, *n* = 34 (siControl) and 27 (siBNIP-2) cells. Median ± interquartile. **d**, Polynomial linear fitting curve of traction force and total area of FAs in siControl (blue curve) and siBNIP-2 (red curve) H9c2 cells. Each data point presents individual cells in four independent experiments. *n* = 19 (siControl) and 18 (siBNIP-2) cells. (**a**,**c**) Unpaired two-tailed Student’s t-test; \**P*<0.05, \*\**P*<0.01, \*\*\**P*<0.001, and \*\*\*\**P*<0.0001.

To investigate how BNIP-2 regulates force transmission via FAs, H9c2 cells transfected with GFP-paxillin and short interfering RNA (siRNA) targeting BNIP-2 were seeded on PDMS gel embedded with fluorescent beads for traction forces measurement (Figure 1b). BNIP-2 knockdown cells showed a significant reduction of traction forces compared to the control cells (Figure 1c). The efficiency of siRNA targeting BNIP-2 and its various isoforms was validated by Western blotting (Figure S1c, Supporting Information). The reduction of traction forces accompanied by a decrease in FA areas is typically observed in fibroblastic cell model^36,41^. However, the morphometric analyses of FAs revealed that BNIP-2 depletion increased the FA mean area per cell while reducing the total FA number, although there was no significant difference in the total FA area per cell compared to control cells (Figure 1c and **Figure S2**a, Supporting Information). Although the cell area is a known confounding factor of FA size, there was no significant change in the cell sizes after BNIP-2 knockdown (Figure S2b-c, Supporting Information). Instead, BNIP-2 knockdown cells revealed a higher proportion of larger FAs (Figure S2d, Supporting Information). To understand the correlation between reduced traction force and larger FAs in BNIP-2 knockdown cells, the total FA area per cell versus traction force per cell was plotted and fitted using a linear regression model (Figure 1d). These results indicate that BNIP-2 knockdown in H9c2 cells led to lower traction forces compared to control cells. Notably, with increasing total FA area per cell, the traction force exerted by BNIP-2 knockdown cells showed a less sensitive response (as denoted by the gradient of the linear fit curve), whereas the control cells have a steeper force increment with the same increment of total FA area compared to the BNIP-2-depleted cells. These results indicated that BNIP-2 knockdown impaired the ability of cells to exert traction force through their FAs and attenuated the coupling between FA force transmission and FA maturation.

Next, H9c2 cells were treated with ROCK inhibitor Y-27632, which is known to deregulate traction forces and FA areas in multiple cell types, to identify if the observation of larger traction forces with smaller FAs is a BNIP-2-dependent or force-regulated phenotype (as shown by BNIP-2 expressing H9c2 in Figure 1b-c)^42, 43^. However, the FA size of H9c2 were decreased after treatment with Y-27632 (**Figure S3**a-b, Supporting Information). These results pointed out the convergent roles BNIP-2 and Rho/ROCK in governing the size of FAs in general. Furthermore, it implies that decreased BNIP-2 expression which resulted in larger FA and diminished contractility, is independent of Rho/ROCK signaling regulation. Instead, BNIP-2 acts as a predominant regulator on the FA size.

### BNIP-2 regulates mobility of FAK, paxillin and vinculin

Since the traction force and FA maturation are attenuated significantly in BNIP-2 KD cells, we hypothesize that BNIP-2 participates in the regulation of FA dynamics. We first addressed the assembly and disassembly rates of FAs using time-lapse imaging (**Figure S4**a-b, Supporting Information). Paxillin and vinculin are used as the FA markers as paxillin is the early and persistent component in FAs, and vinculin, mainly recruited in a force-dependent manner, is involved in the more mature stage in FA formation^44^. After BNIP-2 knockdown, there was a reduction in paxillin assembly and disassembly rates and vinculin assembly rates, and there was no significant change in vinculin disassembly rates. (**Figure 2**a and Figure S4a-c, Supporting Information). The protein mobility of FA proteins was measured in live cells expressing mApple-paxillin, mCherry-vinculin, or mApple-FAK using fluorescence recovery after photobleaching (FRAP) microscopy (Figure 2b-c and Figure S4d, Supporting Information). The results showed that FA proteins differed in their protein-exchange rates within FAs and these rates were significantly affected by BNIP-2 knockdown. The recovery kinetics of FAK [*T*_1/2_ (Control)= 8.2 s, and *T*_1/2_(KD)= 3.4 s] and paxillin [*T*_1/2_(Control)= 6.9 s, and *T*_1/2_ (KD)= 4.0 s] were significantly faster in BNIP-2 knockdown H9c2 cells, whereas the vinculin recovery kinetic was slower after BNIP-2 knockdown [*T*_1/2_ (Control)= 4.3 s, and *T*_1/2_ (KD)= 6.5 s] (Figure 2d). The recovery extent of paxillin was enhanced in BNIP-2 knockdown H9c2 cells, but the vinculin recovery extent was reduced after BNIP-2 knockdown. However, the overall recovery of FAK did not differ significantly between the control and BNIP-2 knockdown H9c2 cells.

**Figure 2.**
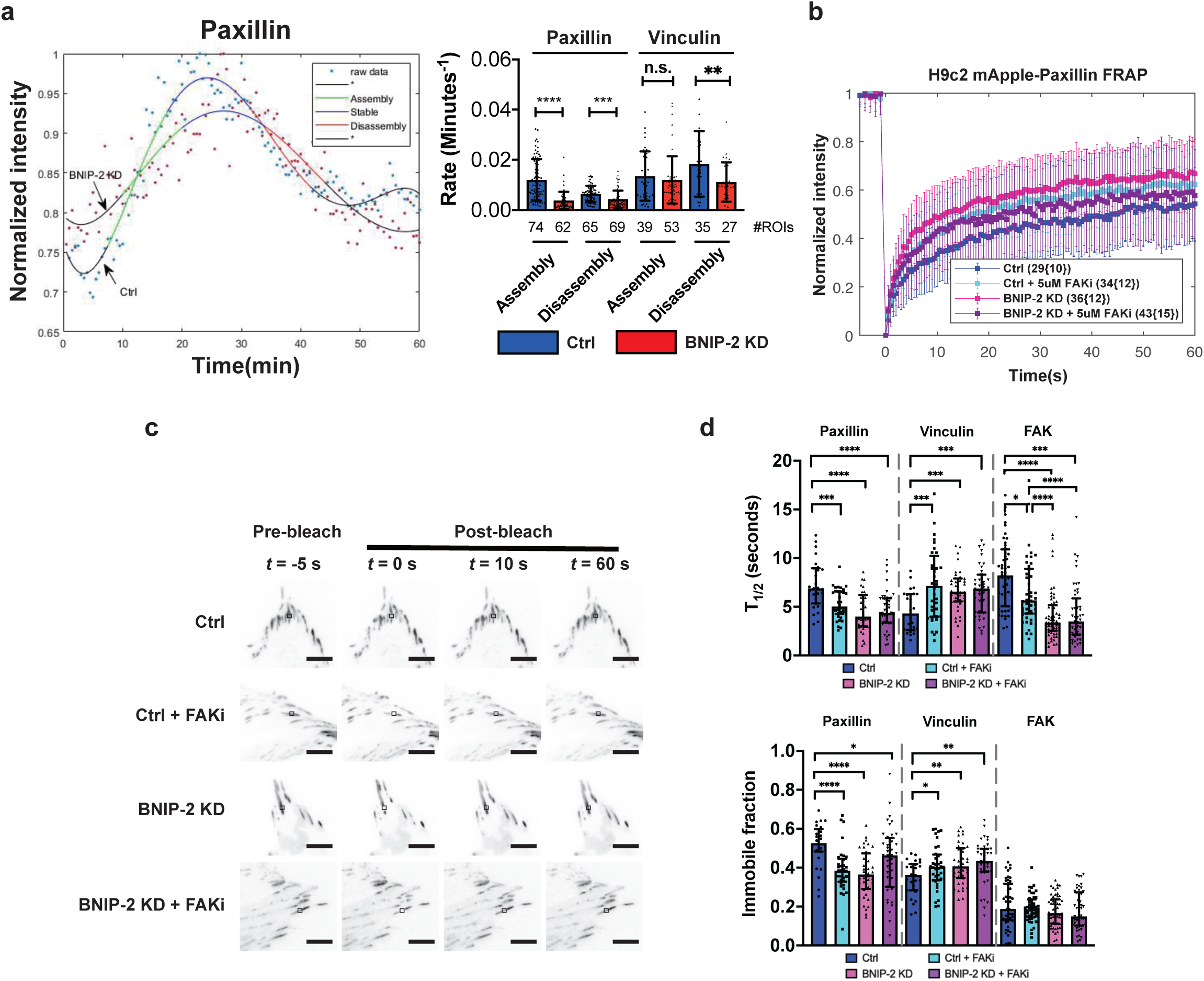
BNIP-2 regulates the mobility of paxillin and vinculin. **a**, Calculation of FA assembly and disassembly rates in H9c2 cells expressing mApple-paxillin and siRNA-targeting BNIP-2 (left). Quantitative analyses of assembly and disassembly rates of paxillin and vinculin (right). **b**, FRAP-relative fluorescence intensity curves of control, BNIP-2-knockdown, and FAK-inhibited H9c2 cells expressing mApple-paxillin. The fluorescence intensity is normalized by the average prebleach intensity, mean ± s.d.. The number of FRAP regions is presented as (number of FAs {number of cells}). **c**, FRAP images of H9c2 cells expressing mApple-paxillin under control, BNIP-2 knockdown, and FAK inhibition conditions. Scale bar, 10 μm. **d**, Calculation of median ± interquartile half-life time (*T*_1/2_) and immobile fraction of each FAs in control, BNIP-2-knockdown, and FAK-inhibited H9c2 cells expressing mApple-paxillin, mCherry-vinculin, and mApple-FAK. Median level of *T*_1/2_ is used as the representative half-life time of each given FA protein (**a**,**d**) Median ± interquartile; Unpaired two-tailed Student’s t-test; \**P*<0.05, \*\**P*<0.01, \*\*\**P*<0.001, and \*\*\*\**P*<0.0001.

Paxillin and FAK contribute to mechanosignaling within FAs whereas vinculin is the force-dependent component involved in mechanosensing^45, 46^. FRAP results reveal that FAK and paxillin molecules are more mobile after BNIP-2 knockdown (Figure 2d). Vinculin, however, is relatively immobile with slower exchange kinetic in BNIP-2 knockdown conditions. The differential dynamics of FA proteins presented above suggest that BNIP-2 affects the turnover of force-independent components in FAs and is necessary for paxillin stability. Meanwhile, the reduced vinculin turnover due to the absence of BNIP-2 can be potentially caused by the lower contraction force after BNIP-2 knockdown, and BNIP-2 could be the key driver of high force generation to control the vinculin turnover. FAK is a regulator of integrin-mediated cell adhesions where FAK inhibition enlarges FA areas and lowers the rate of FA disassembly^47, 48^. The effects of BNIP-2 reduction on FA dynamics phenocopied that of FAK inhibition, and BNIP-2 reduction led to a higher FAK exchange rate than that induced by FAK inhibition (Figure 2d). In addition, FAK inhibition had no further impact on the effects brought about by BNIP-2 knockdown (Figure 2d), suggesting that loss of BNIP-2 has a similar impact as FAK inhibition, likely via a convergent mechanism. Collectively, these results support the role of BNIP-2 in mediating FA dynamics and force transmission.

### BNIP-2 integrates FA machinery and enhances FAK/paxillin signaling

Protein-protein interactions among FA components serve to transduce mechanical signals^49^. As the FA dynamic was affected after BNIP-2 depletion, we hypothesized that BNIP-2 integrates the FA complex functionally and mediates its downstream signaling. We first determined the locality of BNIP-2 and its interaction with FA proteins. Proximity ligation assay (PLA) and co-immunoprecipitation (co-IP) assays were conducted to investigate the complex formation between BNIP-2 with paxillin, vinculin and FAK in H9c2 cells (**Figure 3**a-e, **Figure S5a-f**, Supporting Information), while BNIP-2 colocalization with FA proteins in H9c2s is shown in **Figure S6**a-c, Supporting Information. Protein interaction network analyses using STRING^37, 50^ also predicts that BNIP-2 is likely associated with paxillin as the hub in regulating FA signaling (**Figure S7**, Supporting Information).

**Figure 3.**
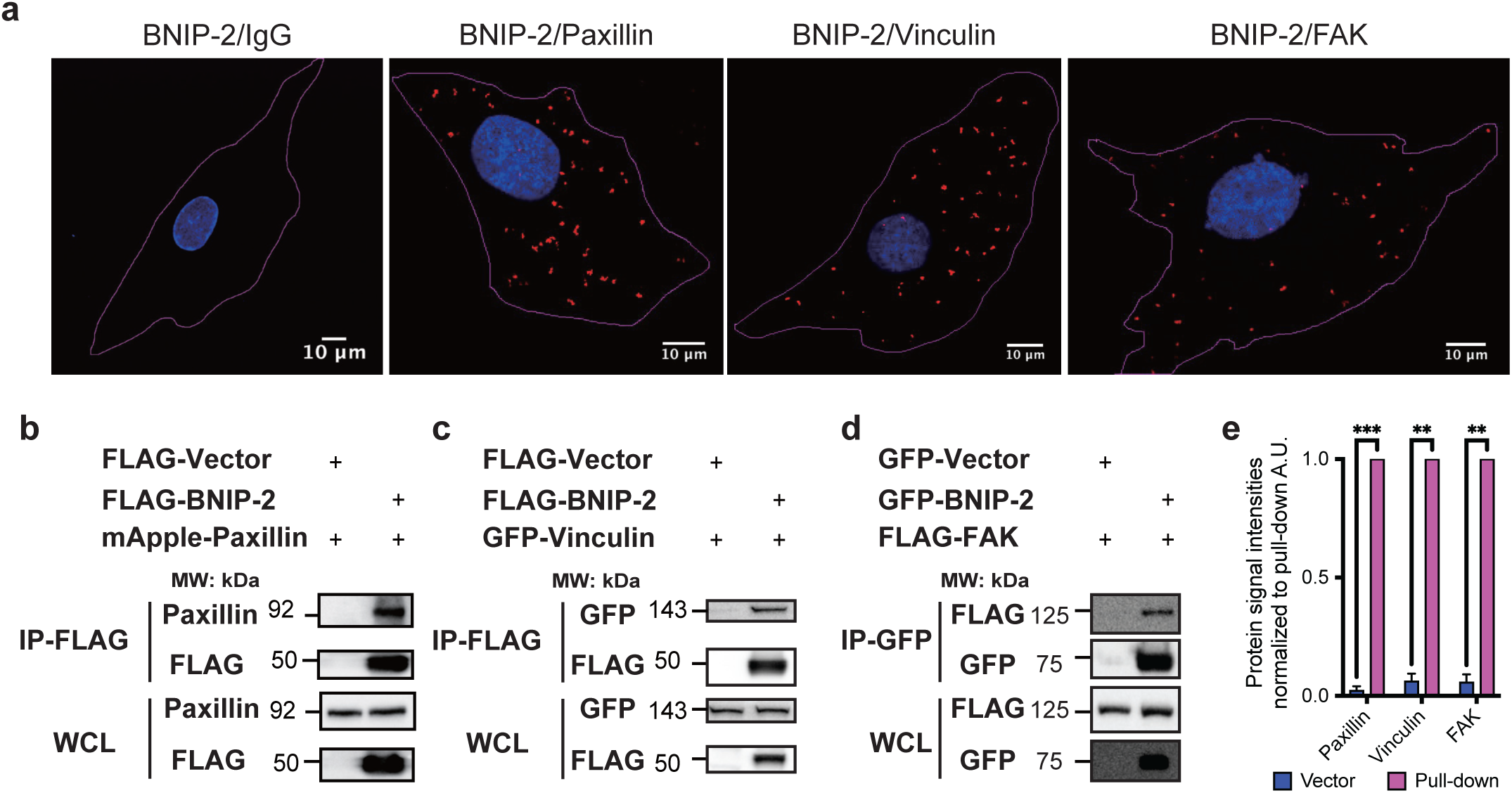
BNIP-2 forms a complex with paxillin, vinculin, and FAK. **a**, Images of interaction detections between BNIP-2 and FA proteins (from left to right, IgG control, paxillin, vinculin, and FAK) in H9c2 myoblasts using proximity ligation assay (PLA). Scale bar, 10 μm. **b**-**d**, Immunoblot analyses of the interaction identification between BNIP-2 and FA proteins (left to right, paxillin, vinculin, and FAK) using co-immunoprecipitation (co-IP) in HEK293T cells expressing mApple-paxillin, GFP-vincullin, FLAG-FAK, and FLAG-tagged or GFP-tagged vector and BNIP-2. Anti-FLAG or anti-GFP magnetic beads were used for each immunoprecipitation according to manufacturer’s instruction. The immunoprecipitation blot probed using anti-paxillin, anti-GFP, or anti-FLAG antibodies indicates the interaction between BNIP-2 and FA proteins (i.e. paxillin, vinculin, FAK). Please refer to Figure S5 for co-IP replicates. **e**, Quantification of immunoprecipitation blots in **b-d** and Figure S5a-f (supporting information) for detecting BNIP-2 interaction with paxillin, vinculin, and FAK. The band intensities were normalized to pull-down lane (the second lane in immunoprecipitation blot). Mean ± s.e.m., n=3 independent experiments. Unpaired two-tailed Student’s t-test; **P<0.01 and ***P<0.001.

FAK phosphorylation of specific tyrosine residues on paxillin promotes the turnover and translocation of FA proteins^51, 52^. Inhibition of FAK/paxillin interactions results in the release of FAK from FAs, thereby reducing the activation of FAK-mediated downstream targets^53^. Combined with the result that BNIP-2 knockdown phenocopied FAK inhibition on FA dynamics regulations, we hypothesized that BNIP-2 promotes force transmission by activating FAK/paxillin signaling. We first examined the activation status of FA proteins in BNIP-2-depleted and FAK-inhibited H9c2 myoblasts. Our data showed that BNIP-2 knockdown phenocopied the effects of FAK inhibition on FA protein phosphorylation (**Figure 4**a and **Figure S8**a, Supporting Information). The quantification of blots (Figure 4b and Figure S8b, Supporting Information) revealed that in both BNIP-2 knockdown and FAK-inhibited H9c2 cells, the level of FAK Y397 and paxillin Y118 were lower compared to the control cells. Notably, the expression level of total paxillin was also lower than in control cells, implying that the loss of BNIP-2 may facilitate paxillin degradation. Contrastingly, the effect of BNIP-2 on vinculin expression and its activation was negligible (Figure 4a-b and Figure S8a-b, Supporting Information). As vinculin serves as a modulator of FAK/paxillin interactions and their phosphorylation to control cell behaviors^20^, this result suggests that vinculin activation is not the key driver of the loss of force transmission after BNIP-2 reduction. FAK Y397 residue is auto-phosphorylated upon FAK interacting with integrin, which provides a high-affinity binding site for Src to recognize and interact^54^. Src interaction with FAK helps stabilize the open conformation of Src kinase and promotes the subsequent FAK phosphorylation at Y576, Y577, and Y925 to promote cell motility, growth, and survival^55^. BNIP-2 knockdown phenocopied FAK inhibition on reduced FAK phosphorylation at Y925, expected from the reduced phosphorylation of FAK Y397 after BNIP-2 depletion (Figure S8c-d, Supporting Information). However, neither BNIP-2 knockdown nor FAK inhibition had significant effects on the expression of Src and phosphorylated Src Y416 in H9c2 cells, highlighting the specificity of BNIP-2 effect on FAK/paxillin signaling (Figure S8c-d, Supporting Information).

**Figure 4.**
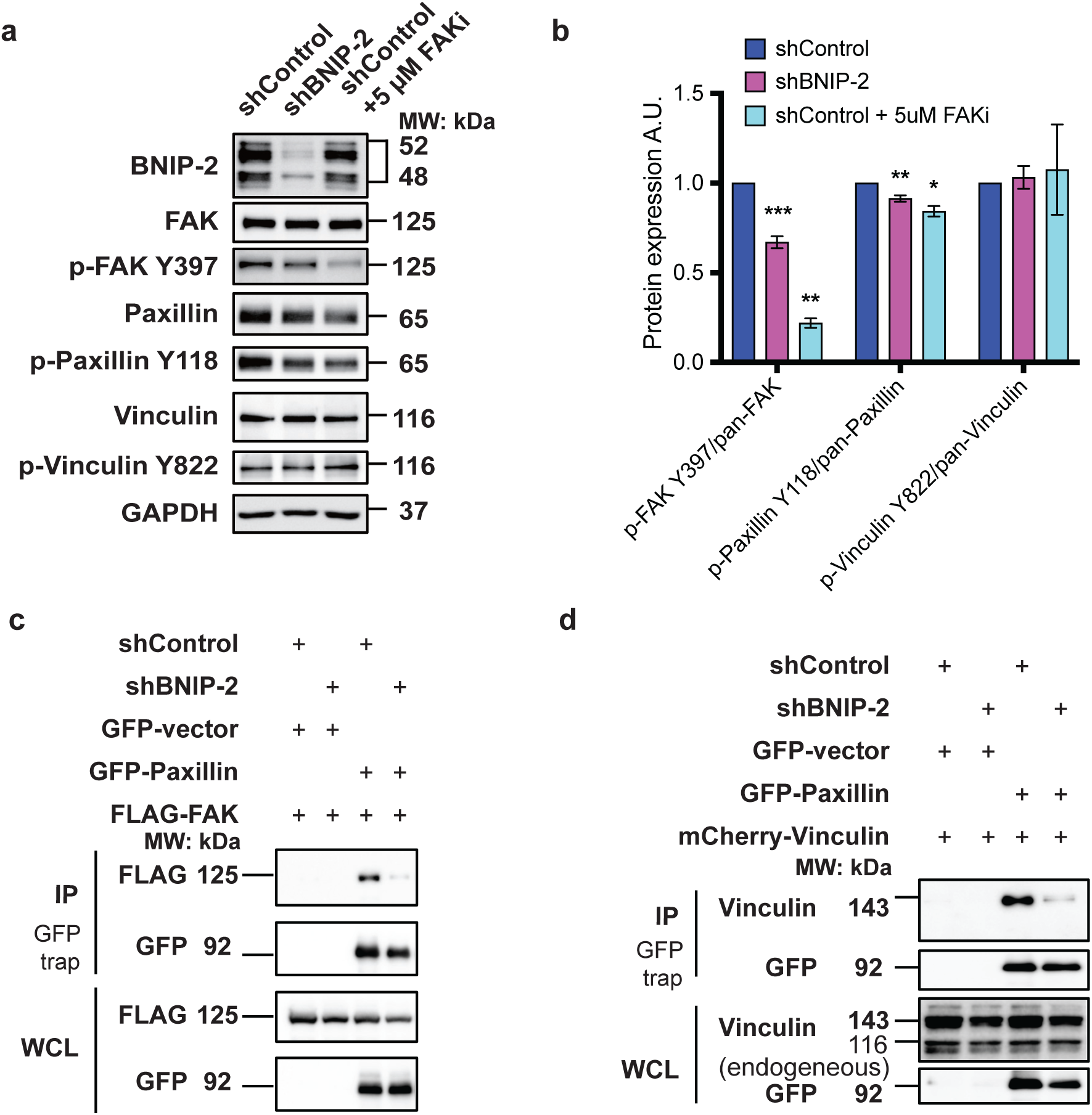
BNIP-2 as a scaffold for FAK/paxillin and paxillin/vinculin interactions. **a**, Immunoblot analyses of BNIP-2, FAK, paxillin, vinculin, phosphorylated FAK Y397, p-paxillin Y118, p-vinculin Y822 in control and BNIP-2 knockdown H9c2 myoblasts. Blots of control H9c2 cells treated with FAK inhibitor are shown. **b**, The level of phosphorylation of each given protein relative to control cells without FAK inhibition. The phosphorylated level was normalized to the total protein expression of each given protein. Mean ± s.e.m., *n*=3 independent experiments. Unpaired two-tailed Student’s t-test; \**P*<0.05, \*\**P*<0.01, and \*\*\**P*<0.001. **c-d**, BNIP-2 knockdown reduces interactions between paxillin and FAK/vinculin in HEK293T cells expressing GFP-tagged paxillin or empty vector and FLAG-tagged FAK or mCherry-vinculin. Immunoprecipitation and immunoblot detection of FAK or vinculin with anti-GFP-trap magnetic agarose beads are shown. The immunoprecipitation blot probed using anti-FLAG or anti-vinculin antibodies indicates the interaction between paxillin and FAK or vinculin, respectively.

Next, we examined the effect of BNIP-2 on the interactions among these FA proteins. Our results showed that less FAK was pulled down by GFP-tagged paxillin in BNIP-2 knockdown cells (Figure 4c). Vinculin interacts with paxillin through its C-terminal within the FAs^56^. The co-IP result also presented reduced interactions between paxillin and vinculin after BNIP-2 depletion (Figure 4d), which may also result from the reduced paxillin expression after BNIP-2 knockdown. The reduction in interactions among FAK, paxillin, and vinculin after BNIP-2 knockdown implies that BNIP-2 serves as a bridge that scaffolds FAK/paxillin/vinculin, consistent with the observed perturbation by reduced BNIP-2 on the exchange mobility of these proteins seen by FRAP. This reduction in the interactions among FA proteins could lead to a decrease in the phosphorylation of several FA proteins, which implicates the mediation in force transmission from the extracellular environment to cytoskeleton organization^57^.

### BNIP-2 as an organizer of FAK localization at ISL

Multiple molecular components form an integrated multi-layered structure within FAs^12^. The distribution of molecular components of FA proteins within FAs is essential for its function in mechanotransduction. To further examine how BNIP-2 scaffolds the FAK/paxillin/vinculin complex, Scanning angle interference microscopy (SAIM) was used to map the organization of FA proteins at the nanoscale^58, 59^. To determine how BNIP-2 contributes to force transmission regulation by modulating FA structure organization, H9c2 cells expressing siRNA targeting BNIP-2 and fluorophore-tagged FA proteins were imaged and measured for their Z-position after fixation (**Figure 5**a and **Figure S9**a, Supporting Information). Figure 5b and Figure S9b show the histogram and box plots of the FA proteins distribution in Z. The median Z-level of FA proteins is used as the representative Z-level for each given FA protein, depicted in the schematic diagram of FA nanoscale architecture in H9c2 cells (Figure S9c, Supporting Information).

**Figure 5.**
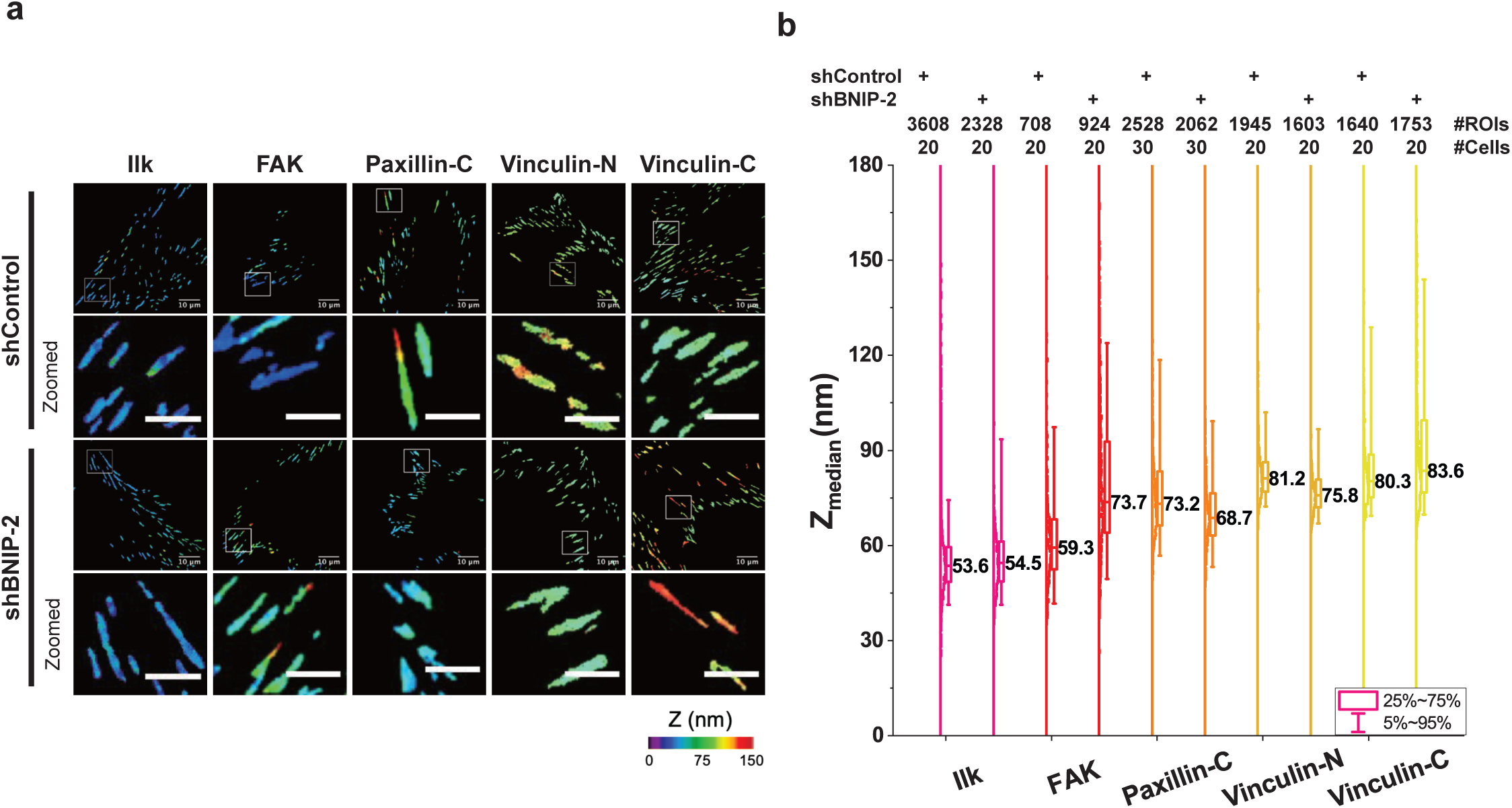
BNIP-2 as an organizer of FAK localization at integrin signaling layer. **a**, Color-coded topographic map in Z-axis of FA proteins in control and BNIP-2 knockdown H9c2 cells expressing fluorophore-tagged FA proteins (i.e. ILK, FAK, paxillin-C, vinculin-N, and vinculin-C). Color bar indicates the Z-position relative to the surface. Scale bar, 10 μm (original) and 5 μm (zoomed). **b**, Histogram and box plot of FA Z-positions. Each data point represents the Z-center of individual FA region of interest (ROI) relative to the surface. The number of ROIs and cells labelled at the top of the plot. Median ± interquartile.

Integrin-linked kinase (ILK) is used to demarcate the plasma membrane Z-level, and both control and BNIP-2 knockdown H9c2 had the ILK at a similar Z level (Z_Control_ = 53.6 nm, and Z_KD_ = 54.5 nm). The organization of other major components of FAs, including talin, tensin, vinculin, and zyxin, were unaltered by BNIP-2 depletion, indicating the force transduction layer and actin regulatory layer were not perturbed by BNIP-2 knockdown (Figure S9a-b, Supporting Information). However, the ISL component of FAK was observed to be at a much higher Z level in the BNIP-2 knockdown condition, Z_Control_ = 59.3 nm vs Z_KD_ = 73.7 nm, whereas the changes of paxillin Z-level were much less than that of FAK after BNIP-2 depletion (Z_Control_ = 73.2 nm, and Z_KD_ = 68.7 nm) (Figure 5a-b). Therefore, BNIP-2 modulates the organization of ISL in FA structure where its depletion significantly displaced the Z position of FAK with respect to other components of FAs. This result explained the reduced FAK/paxillin interactions and FAK-mediated signaling in BNIP-2-depleted H9c2 cells. As the FAK inhibition phenocopied the BNIP-2-regulated FA dynamics and signaling, the reorganized FAK spatial distribution implied that the BNIP-2 regulates FAK/paxillin signaling and force transmission through mediating FAK distribution within FAs.

### BNIP-2-BCH domain tethers the interaction between FAK and BNIP-2 to modulate Paxillin phosphorylation

FAK autophosphorylation site Y397 is indispensable for FAK function within FAs^55^. The inhibition of FAK using PF-562271 demonstrates a higher proportion of FAK localized in the nucleus^60^. As BNIP-2 knockdown phenocopied FAK inhibition on regulating the activation status of FA proteins, we wondered if FAK phosphomimetic mutants could rescue the effect of BNIP-2 knockdown on FA signaling. We first compared the interactions between BNIP-2 and phosphomimetic FAK mutants. FAK Y397D showed enhanced interaction with BNIP-2, whereas the FAK Y397F barely interacted with BNIP-2 (**Figure 6**a), suggesting that BNIP-2 forms a more stable complex with activated FAK. Myosin light chain (MLC) and its phosphorylation serve to regulate cell contraction^61, 62^. Interestingly, overexpressing FAK Y397D, but not wildtype FAK, was able to rescue paxillin phosphorylation and contractility, as noted by the increased phosphorylation of paxillin Y118 and MLC T18/S19 (Figure 6b). The ability of FAK Y397D to bypass and rescue the loss of paxillin and MLC activation due to BNIP-2 depletion suggests that FAK/paxillin/MLC activation occurs downstream of BNIP-2.

**Figure 6.**
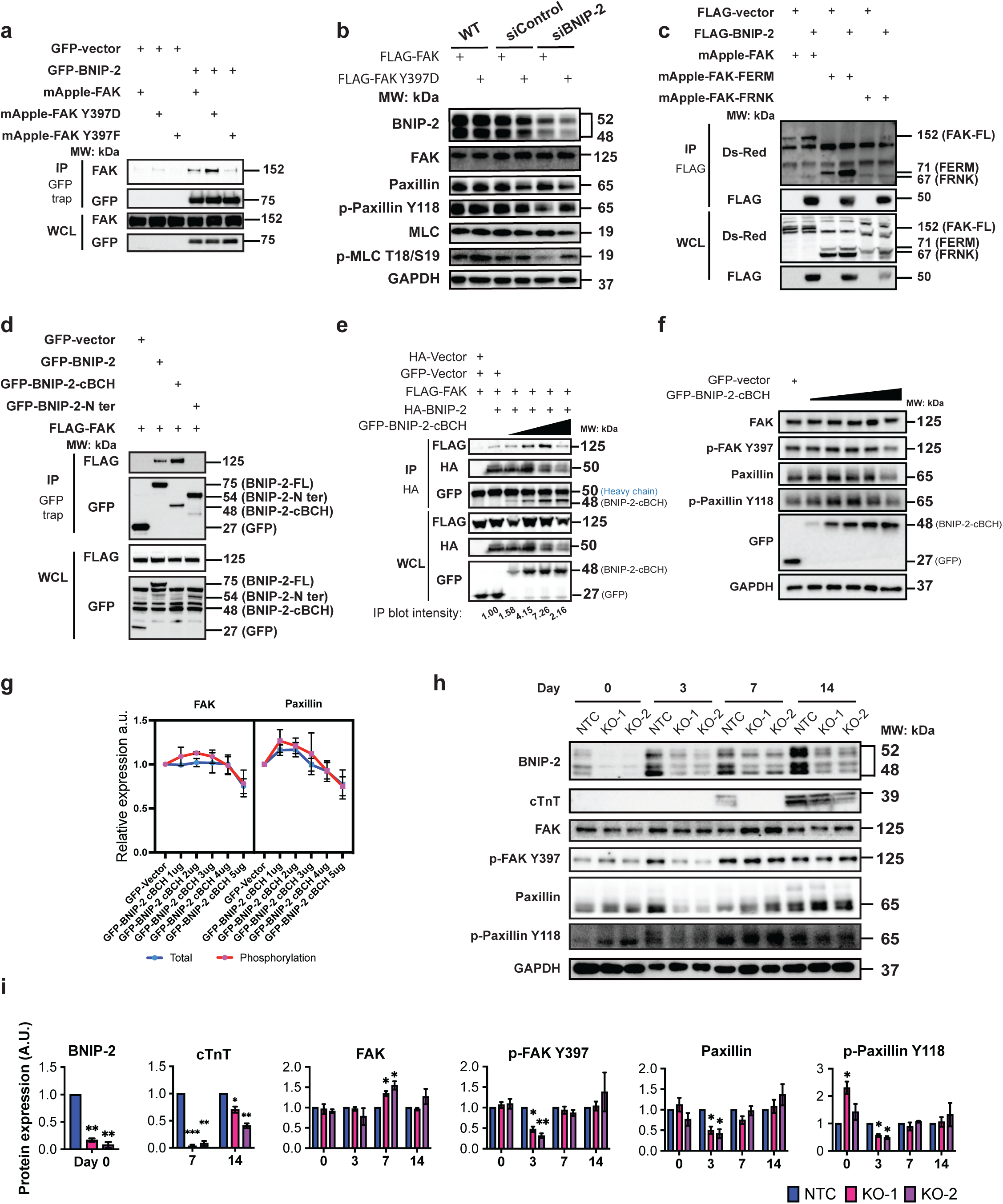
BNIP-2 interacts with FAK FERM domain through its BCH domain to regulate hESC cardiomyocyte differentiation. **a**, Immunoprecipitation analyses of the interaction between BNIP-2 and FAK phosphomimetic mutants in HEK 293T cells expressing GFP-tagged BNIP-2 or empty vector and mApple-tagged FAK Y397D (tyrosine-phosphorylated form) and FAK Y397F (non-phosphorylated form). Immunoprecipitation blot probed using anti-FAK antibody indicates the interaction between BNIP-2 and FAK mutants. **b**, Immunoblot analyses of BNIP-2, FAK, paxillin, phosphorylated paxillin Y118, myosin light chain (MLC), and phosphorylated MLC T18/S19 in H9c2 cells transfected with siRNA-targeted BNIP-2, FLAG-FAK, and FLAG-FAK Y397D respectively. **c**, Immunoprecipitation of FAK truncated mutants and BNIP-2 in HEK293T cells expressing GFP-tagged empty vector or BNIP-2 and mApple-tagged FAK truncated mutants (FAK-FL: FAK full-length; FERM: FAK-FERM domain on the N-terminal; FRNK: FAK-FRNK domain on the C-terminal; please refer to Figure S10b for illustration of regions, Supporting Information). **d**, Immunoprecipitation analyses of the interaction between BNIP-2 truncated mutants and FAK in HEK293T cells expressing GFP-tagged BNIP-2 truncated mutants or empty vector and FLAG-tagged FAK (please refer to Figure S10c for illustration of regions, Supporting Information). **e**, Immunoprecipitation analyses of BNIP-2-FAK interactions in HEK293T cells expressing increasing amount of GFP-BNIP-2cBCH (illustrated by black triangle). Anti-HA magnetic beads were used according to manufacturer’s instruction. Heavy chain of anti-HA beads is labelled. The immunoprecipitation blot intensities were quantified from the anti-FLAG blot in IP sample and normalized to the anti-HA blot in IP sample, which represents the total pull-down amount. **f**-**g**, Immunoblot analyses of FAK, phosphorylated FAK Y397, paxillin, phosphorylated paxillin Y118, and GFP in H9c2 myoblasts expressing gradient amount of GFP-BNIP-2-cBCH (illustrated by black triangle). Anti-GFP blot denotes the expression of GFP-BNIP-2-cBCH. **h**-**i**, Immunoblot analyses of BNIP-2, cTnT, FAK, paxillin, phosphorylated FAK Y397 and p-paxillin Y118 in non-targeting control (NTC), CRISPR-targeted BNIP-2 knockdown hESCs on Day 0, 3, 7, and 14 of cardiomyocyte differentiation. Non_Targeting_sgRNA1 and Non_Targeting_sgRNA2 were applied for control hESC stable cells generation (NTC); BNIP-2_sgRNA1/BNIP-2_sgRNA3 and BNIP-2_sgRNA2/BNIP-2_sgRNA4) were used for generating BNIP-2 knockdown hESC KO-1 and KO-2 separately. Mean ± s.e.m.; Unpaired two-tailed Student’s t-test; \**P*<0.05, \*\**P*<0.01, and \*\*\**P*<0.001. (**a**-**f**, **h)** *n*=3 independent experiments.

FAK consists of a FERM domain on the N-terminal, a central catalytic kinase domain, and a focal adhesion targeting (FAT) on the C-terminal^63^. Truncated FAK mutants were used to investigate the binding domain of FAK to BNIP-2 (**Figure S10**a, Supporting Information). Both full-length FAK and FAK-FERM domain (with the linker between FERM and kinase domain) were pulled down by GFP-BNIP-2 (Figure 6c), indicating that FAK interacts with BNIP-2 through its FERM domain. BNIP-2 is a scaffold protein comprising a functional BCH domain and an unstructured N-terminal domain^32^. To study the functional domain of BNIP-2 interacting with FAK, BNIP-2 was truncated into BCH domain-containing part (BNIP-2-cBCH), and N-terminal domain-containing BNIP-2 (BNIP-2-N) (Figure S10b, Supporting Information). BNIP-2-cBCH contains a short region of N terminal BNIP-2 to improve the stability of the protein expression. The result from co-IP revealed that both full-length BNIP-2 and BNIP-2-cBCH interacted with FAK, while the BNIP-2-N terminus was unable to form a complex with FAK (Figure 6d). BNIP-2, therefore, bridged the FA proteins together to mediate FAK signaling through the BNIP-2-BCH and FAK-FERM domain. The active form of FAK, which mimics the FAK activation status in the presence of BNIP-2, can constitutively bypass and drive the FAK/paxillin signalling and force transmission without BNIP-2. We further showed that BNIP-2-cBCH domain exhibit a critical concentration-dependent scaffolding effect for BNIP-2 and FAK interactions (Figure 6e). Increasing overexpression of BNIP-2-cBCH domain enhanced the BNIP-2/FAK complex formation in a concentration-dependent manner. However, when the level of BNIP-2-cBCH expression went beyond the optimal concentration, it could effectively compete with FAK to interact with full-length BNIP-2 and, therefore, disrupted FAK/BNIP-2 interactions and exhibited the classical scaffolding protein behaviour (Figure 6e). Consequently, the scaffolding effect of BNIP-2-cBCH domain was also involved in tethering FAK/paxillin signaling (Figure 6f-g, **Figure S11**a-b, Supporting Information). Here, the levels of phosphorylated FAK Y397, and phosphorylated paxillin Y118 were enhanced first and were reduced afterward, especially with a much higher amount of BNIP-2-cBCH. In addition, the total paxillin level was also reduced in the presence of high amount of BNIP-2-cBCH domain, suggestive of a functional BNIP-2/FAK/paxillin interaction to maintain paxillin stability. In summary, these results demonstrate that the BNIP-2-cBCH is the domain that allows BNIP-2 to serve as a scaffold for regulating FAK/paxillin signalling. Notably, high amount of BNIP-2-cBCH domain acts as a dominant negative mutant that disrupts their interactions and phenocopies the effect of BNIP-2 knockdown and FAK inhibition on FAK/paxillin signaling.

### Mechanotransduction through BNIP-2-regulated FAK signaling is essential for hESCs cardiomyocyte differentiation

Human embryonic stem cells (hESCs) represents a valuable cell model for studying mechanotransduction as their differentiation status is strongly associated with the extracellular stiffness and force transmission within the cytoskeleton^64^. We have performed CRISPR-targeted BNIP-2 in hESCs to investigate whether BNIP-2 mediates the cardiomyocyte differentiation from hESCs through its regulation of FAK signaling (Figure 6h-i). Loss of BNIP-2 has impeded the hESCs differentiation into cardiomyocytes, as noted in the absence of the cardiomyocyte maturation marker cTnT in BNIP-2-deleted hESCs on Day 7, and cTnT was moderately expressed on Day 14 compared to non-targeting control (NTC) hESCs. Similar to the regulatory role of BNIP-2 on undifferentiated H9c2 cells (Figure 4a-b), FAK/paxillin signaling was inhibited in differentiated BNIP-2-KO hESCs on Day 3 (before cardiomyocyte maturation), during the stage of cardiac mesodermal cell differentiation ^65, 66^. The expression of paxillin, phosphorylated FAK Y397, and phosphorylated paxillin Y118 was barely detected on Day 3 in BNIP-2 knockout hESCs. These results suggest that BNIP-2 primes and directs hESC differentiation by regulating FAK-mediated mechano-signaling in the cardiac mesodermal stage, which is crucial for cardiomyocyte maturation. Intriguingly, BNIP-2 KO hESCs had higher total FAK expression at the matured stage (Day 7). These could be due to the feedback compensatory mechanism for the reduced phosphorylated FAK Y397 at the mesodermal stage, and this compensation was lost during the continued culturing of the matured cardiomyocytes (Day 14). Taken together, our results showed that BNIP-2-regulated spatiotemporal FAK signaling is essential for directing hESCs cardiomyocyte differentiation.

## Discussion

FAs play crucial roles in regulating force transmission and cell differentiation^18^. Here, we have identified BNIP-2 as a novel regulator of FA nanostructure and mechano-signaling, which regulates force transmission in H9c2 myoblasts and mediates hESCs cardiomyocyte differentiation. Mechanistically, BNIP-2 is functionally located at FA, where it play as a spatiotemporal regulator of mechanotransduction by coordinating FA structure and maintaining interactions among FA proteins to elicit force transmission (**Figure S12,** Supporting Information). The 3D mapping of various FA proteins and domain binding studies reveal how BNIP-2 controls the Z-positions of FAK, a vital component of the integrin signaling layer, thereby providing a structural basis for a potential mechanism of how BNIP-2 regulates FAK/paxillin interactions. Our result demonstrated that the reduction of BNIP-2 disrupts the interactions of FAK/paxillin and paxillin/vinculin and decreases the phosphorylation of FAK and paxillin without changing the vinculin expression and activation. The displacement of FAK Z-position away from ISL is first identified in cells with low traction force (BNIP-2 knockdown H9c2). We have also shown that BNIP-2 interacts with the FAK-FERM domain through its BCH domain. These results demonstrate that the BNIP-2 regulation on FAK activity through the FAK-FERM domain plays a key role in mediating FAK downstream signaling and force transmission.

BNIP-2 is known to interact with Rho and GEF-H1 to mediate microtubule-linked actin filament polymerization and force transmission^30, 33^. Reduced BNIP-2 expression lowered the Rho activity and downregulated the force transmission, resulting in lower traction forces and fewer activated FA proteins. Here, BNIP-2 regulation leads to force transmission mediated through Rho-dependent and FAK/paxillin signaling. The cell contractility is positively correlated to the FA sizes^41^. However, the reduction of BNIP-2 perturbed force transmission and promoted the increase of FA sizes instead. The assembly measurement of FAs showed a decrease in paxillin disassembly and assembly rates after BNIP-2 reduction (Figure 2a). Likewise, the vinculin and Rho-dependent myosin light chain phosphorylation is required for the FA assembly^24, 69^, and reduced Rho activity decreases FA formation. Additionally, the BNIP-2 at FAs modulates FAK phosphorylation and promotes the paxillin disassembly from FAs. The lack of BNIP-2 leads to a reduced FAK phosphorylation level, and less paxillin is activated at Y118 resulting in paxillin disassembly at FAs. Therefore, the larger FAs could arise from the compensatory result of FA assembly and disassembly influenced by BNIP-2 depletion.

FAK inhibition is known to reduce FA disassembly and increase FA size^47, 48^. Our data show that the loss of BNIP-2 phenocopies FAK inhibition to reduce FAK activation at Y397 and increases the area within individual FA, despite maintaining the total FA area per cell. This phenocopy implies the convergent regulatory role of BNIP-2 on FAK to promote FAK signaling and its downstream force transmission. Meanwhile, BNIP-2 knockdown reduces phosphorylated FAK Y397. However, FAK Y397 is an autophosphorylation site. As FAK is recruited to FAs through its interaction with paxillin, and this interaction could promote the FAK phosphorylation at Y397^70, 71^, BNIP-2 regulates the FAK phosphorylation at Y397 might result from the reduced paxillin/FAK interactions. Additionally, BNIP-2 may act as an anchor for p-FAK Y397 (since BNIP-2 binds FAK-Y397D much more strongly than FAK-Y397F) and prevent it from being dephosphorylated to ensure force-transmitting FA structure and subsequent downstream signaling. Alternatively, BNIP-2 may indirectly recruit other proteins to initiate the activation cascade.

The FA 3D structure demonstrates the distance among FA proteins in Z-axis. This mapping reveals the layering of protein-protein interactions and signal transduction within FAs. Inhibition of FAK/paxillin interactions results in the release of FAK from FAs and reduces phosphorylation of FAK and FAK-mediated downstream targets^53^. However, it is not clear why the turnover rate of paxillin is higher after BNIP-2 knockdown, regardless of its reduced disassembly rate and phosphorylation. It has been proposed that phosphorylated paxillin recruits inactive vinculin to FAs and induces the further activation of vinculin through vinculin-binding partners, including talin and actin^72^. In our study, the reduced paxillin/vinculin interaction and less phosphorylated paxillin after BNIP-2 knockdown do not affect vinculin phosphorylation. The unaffected vinculin phosphorylation suggests that phosphorylation alone is not the key driver of subsequent force transmission. Vinculin phosphomimetic mutants are known to modulate the interactions between paxillin and FAK as well as their phosphorylation^20^, and our results also demonstrate it is not a mutual effect as the loss of paxillin/FAK interactions does not regulate vinculin phosphorylation. Instead, reduced traction forces after BNIP-2 knockdown suggests the interaction among FA proteins and the FA spatial organization are essential for regulating force transmission. Hence, we propose that BNIP-2 may serve to scaffold vinculin and paxillin together to connect and therefore consolidate the FA nanostructure for force transmission. A recent study shows an increased paxillin/vinculin interaction occurs within focal complexes^73^, suggesting that vinculin may serve as an anchor to interact with and stabilize paxillin molecules to promote FA formation. These could explain the higher turnover rate of paxillin in BNIP-2 knockdown H9c2 cells.

Taking these observations together, BNIP-2 is actively involved in force transmission through its regulation of both RhoA activity and FAK-mediated signaling. In hESC differentiation study, the BNIP-2-deleted cells exhibited a delay in cardiac differentiation as noted by the cTnT expression on Day 14 (Figure 6h), thus strongly suggesting that the loss of BNIP-2 did not completely hinder the cardiac differentiation. Instead, cells may possess an inherent compensatory mechanism to counteract the deleterious effect of BNIP-2 deletion and restore their vital physiological function. Current work is on-going to delineate how both arms of Rho and FAK signaling are further coordinated by BNIP-2 in cardiac differentiation and in other tissue growth and development. The Hippo-YAP acts as a tumor suppressor and regulates tissue growth, and YAP serves as a mechano-transducer, where its localization and activation are strongly associated with force transmission^74–76^. Recently, we have shown that BNIP-2 regulation of Hippo-YAP in H9c2 myoblasts through controlling YAP retention in the cytosol^29^. The BNIP-2 may play a key role in FAK-Rho-YAP signaling to control cell migration, proliferation, and differentiation.

In summary, using H9c2 and hESCs models and interrogating the functional interaction of BNIP-2 with FA complex and their mutants with TFM, FRAP, SAIM and protein-protein interaction studies, we have uncovered a novel mechanotransduction hub at FAs that drives force-dependent cardiomyocyte differentiation. Mechanistically, BNIP-2 coordinates the integration of FAK, paxillin and vinculin at FAs and the local spatial distribution of FAK, leading to the activation of FAK on paxillin signaling to ensure downstream force transmission. As the intricacy of FA signaling can be harnessed to enhance the differentiation of cardiomyocytes, this novel mechanism can offer new understanding and future applications to heart regenerative medicine. Since integrin-mediated adhesions play a regulatory role in diseased hearts^9^, BNIP-2-scaffolded FAK/Paxillin signaling in force transmission can also serve as a potential therapeutic target to prevent and manage force-dependent heart diseases.

## Materials and Methods

### Cell culture and chemicals

H9c2 myoblasts from the American Type Culture Collection (ATCC) were cultured in DMEM medium (Hyclone) supplemented with 10% FBS (Gibco) and 1% penicillin-streptomycin (Hyclone), and 1mM sodium pyruvate (Gibco) under 5% CO2 at 37℃. H9c2 were sub-cultured when reaching 70-80% confluence and were plated at 100% confluence before initiating differentiation. 1$M all-trans retinoic acid (ATRA) in low serum DMEM medium (1% FBS) was performed for H9c2 cardiac differentiation. Differentiation medium (1% FBS, DMEM) with the fresh addition of ATRA was replaced daily in the dark for three days. Cells were treated with 5uM FAK inhibitor PF-573228 (Sigma-Aldrich) and 40uM ROCK inhibitor Y-27632 (Sigma-Aldrich) in culture medium (10% FBS) or differentiation medium (1% FBS) accordingly. HEK293T cells were cultured in RPMI 1640 medium (Hyclone) supplemented with 10% FBS (Gibco), 10mM HEPES, and 1% penicillin-streptomycin (Hyclone). All cells were tested negative for mycoplasma.

### Generation of knockdown stable cell line

The pGFP-V-RS vectors (OriGene) expressed BNIP-2 targeting sequences were used for plasmid-based RNAi. The rat BNIP-2 targeting sequence was purchased from Integrated DNA Technologies (IDT) and was applied for generating stable knockdown H9c2 cells (shBNIP-2: 5’-GGAAGGTGTGGAACTGAAAGA-3’). HEK293T BNIP-2 knockdown stable cells were generated as described(1). Western blot was performed for measuring the knockdown efficiency.

### Lentiviral transduction and generation of CRISPR-targeted knockout hESC

sgRNAs were designed and applied for hESC CRISPR-targeted knockout. According to the instructions from website http://crispor.tefor.net/ as described previously(2), the NGG PAM sites targeting exon 1, 3, and 4 of the BNIP-2 gene locus were detected for designing sgRNAs. The “ Rule set 2” scoring model was used for prioritizing top sgRNA candidates(3)

Non_Targeting_sgRNA1: 5’-AAAACAGGACGATGTGCGGC-3’;

Non_Targeting_sgRNA2: 5’-AACGTGCTGACGATGCGGGC-3’;

BNIP2_sgRNA1 (Exon 3): 5’-ACTAGCTATAACTGGACCAG-3’;

BNIP2_sgRNA3 (Exon 1): 5’-GATAACATCCCGACCTCCTC-3’;

BNIP2_sgRNA2 (Exon 4): 5’-GACAATACAGAGCCATCACT-3’;

BNIP2_sgRNA4 (Exon 1): 5’-GGTCTCCACCGCCGACCGAG-3’.

Esp31-digested LentiCRISORv2 backbone (Addgene #52961) was encoded annealed sgRNA constructs using T4 DNA ligase (catalog #: M0202, New England Biolabs, NEB). Constructs were confirmed by Sanger sequencing. Lentiviruses were produced in HEK293T cells by co-transfection with10 µg of sgRNA plasmid construct, 7.5 µg of pMDLg/pRRE, 2.5 µg of pRSV-Rev, and 2.5 µg pMD2.G (Addgene #12251, #12253 & #12259) using 50 µl of PEI and 3ml of Opti-MEM™ I Reduced Serum Medium (catalog #31985070, ThermoFisher Scientific), where cells are cultured on 10cm dish with 70% confluence. The medium was changed to DMEM supplemented with 5% FBS after overnight incubation. The supernatant was collected and filtered twice after 24 hours and 48 hours, then concentrated using Lenti-Pac™ Lentivirus Concentration Solution (catalog #: LPR-LCS-01, GeneCopoeia) before transduction. The hESC cell culture medium was supplemented with 8 µg/ml polybrene for increasing transduction efficiency. Cells were purified further using 1 ug/ml Puromycin (Sigma catalog #: P9620) for two days and sub-cultured for at least 5 days prior to cardiomyocyte differentiation.

### Human embryonic stem cell differentiation assays

Geltrex (ThermoFisher Scientific, catalog #: A1413202) was applied to coat culture vessels for 30 minutes before seeding cells. H1 human embryonic stem-cell (H1-hESC) line was cultured in mTeSR1 medium (STEMCELL Technologies) under 5% CO2 at 37℃ and was sub-cultured when reaching 80% confluence. Cells were treated with Accutase (Invitrogen) and resuspended in media supplemented with 10$M ROCK inhibitor Y27632 (STEMCELL Technologies). Cells were then seeded onto Geltrex-coated plates (12-well) at a density of 200,000 cells/cm^2^. The medium was replaced by the culture medium (without ROCK inhibitor) after 24 hours. The cardiomyocyte differentiation GiWi protocol (GSK3/Wnt inhibitor) with RPMI differentiation medium was adopted as described^65^.

### Immunofluorescence

Cells were fixed with pre-warmed 4% paraformaldehyde in PBS for 15 min at 37℃ and washed for 5 min in 1x PBS three times. Samples were then blocked with 3% bovine serum albumin and 0.1% Triton X-100 in 1X PBS for 30 min. Following the primary antibody incubation for 60 min at room temperature, where primary antibodies were diluted in an appropriate ratio in blocking solution, samples were washed by 1X PBS three times and incubated with Alexa Fluor-conjugated secondary antibodies in the dark for 30 min at room temperature. Three times 1X PBS washes were performed before image acquisition.

### Traction force microscopy

Dishes with PDMS gel for measuring traction force were prepared as described previously^77^, where codes and analysis methods can also be found. H9c2 cells were seeded on 6-well plates in 60% confluence, followed by siRNA and GFP-paxillin transfection. Cells were trypsinized, plated onto glass-bottom dishes with PDMS gel (15 kPa, coated with fibronectin), and incubated overnight prior to image acquisition. Images of fluorescent beads before and after 10% SDS treatment were acquired for traction force analysis. Paxillin signals were recorded for the quantification of focal adhesion sizes. OriginPro 2019b was used for generating the linear fitting curve.

### Confocal microscopy

Confocal images were acquired using Yokogawa CSU-W1 Spinning Disk microscope (Nikon) equipped with a 60X 1.20 NA water immersion objective or 100X 1.45 NA oil immersion objective. Confocal microscopy was performed for the traction force microscopy and the detection of BNIP-2/focal adhesion proteins localization.

### Focal adhesion morphometrics and quantification

Focal adhesion quantification was carried out in ImageJ and MATLAB. “Subtract background” was applied with the rolling ball radius set to 50 pixels, followed by the contrast enhancement and the Gaussian blur with a sigma of 0.5. Particles with an area of between 0.1 and 25 µm^2^ were considered and segmented for further analysis. The area of each focal adhesion was recorded as a region of interest (ROI) in ImageJ, and the number and total area of focal adhesions per cell were calculated using MATLAB.

### Fluorescence recovery after photobleaching (FRAP)

FRAP images were acquired on Olympus IX81 inverted microscope (iLAS2 module) with a 100x 1.49 NA oil immersion objective in Total Internal Reflection Fluorescence (TIRF) mode. FRAP experiments were done at 37℃ with 5% CO2. H9c2 cells expressing mApple-paxillin, mApple-FAK, or mCherry-vinculin were seeded on 27 mm glass-bottom dishes for 24 hours before FRAP experiments. Photobleaching was set up as pre-bleaching for 5 s at 1-s intervals, 600 ms for photobleaching, and 60 s post-bleach imaging at 500-ms intervals. Cells were incubated for 1 hour in the culture medium supplemented with 5 μM FAK inhibitor PF-573228 (Sigma-Aldrich) before FRAP experiments.

### Quantitative analysis of FRAP

Plots were generated using ImageJ and MATLAB. ROIs were manually selected on images, including the unbleached area of focal adhesions for bleach-control, empty area for background measurements, and bleached area of focal adhesions for focal adhesion turnover quantification. The integrated intensity of each ROI was measured in ImageJ, followed by background/photobleaching correction performing the formula

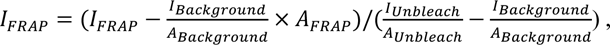

where *I_background_* and *I_unbleach_* were the integrated intensities of background ROI and bleach correction ROI (unbleached ROI), and *A_Background_* and *I_unbleach_* were the area of background ROI and bleach correction ROI, respectively.

FRAP intensities were normalized relative to each FRAP image’s average pre-bleach fluorescence intensity. Following normalization of FRAP intensities relative to the average pre-bleach fluorescence intensity, the normalized intensity data were then performed data fitting to acquire the half-life time *T*_1/2_ via MATLAB using the equation

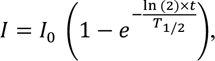

where *I* was the normalized intensity during the FRAP recovery, *I*_0_ was the normalized end-value of recovered intensity (mobile fraction), and t was the time post bleaching.

### Calculation of focal adhesion assembly and disassembly rates

Live imaging was acquired using confocal microscopy (60X objective) as described in the confocal microscopy method. H9c2 cells expressing shRNA and mApple-paxillin/ mCherry-vinculin were seeded on 27 mm glass-bottom dishes for 24 hours before imaging. Images were captured at 37℃ with 5% CO2 for 1 hour at 1-min intervals. Focal adhesions were manually tracked frame by frame in ImageJ, where the integrated intensity was measured. Intensities of each time-lapse image were normalized in MATLAB, followed by the data fitting with a polynomial model of degree 5 to determine the phase lengths of assembly and disassembly. Data points in each phase were then processed with the linear fitting model (3 = 45 + 7) to calculate the rates in the time series.

### Scanning angle interference microscopy (SAIM)

Scanning angle interference microscopy was performed for determining the protein position in the z-axis as previously described^58, 59^, where silicon wafer (Bonda Technology) containing a layer of thermal SiO2 (∼500 nm) was used as the imaging substrate. Following the fibronectin (Sigma-Aldrich, FC010) coating on silicon wafers with 10 8g/mL concentration for 1 hour at 37℃, H9c2 cells were seeded on ECM coated silicon wafers and then transfected with focal adhesion proteins tagged with mApple or mCherry for 24 hours before fixation. Cells were fixed by pre-warmed 4% paraformaldehyde for 15 min at 37℃ and washed with 1X PHEM buffer three times (2 mM MgSO4, 10 mM EGTA, 25 mM HEPES, and 60 mM PIPES; pH 7.0). Images were captured using TIRF microscopy (Nikon Eclipse Ti inverted microscope) equipped with a 60X 1.49 NA objective (Nikon Instruments). Wafers with cells were flipped and facing downward in a 27 mm glass-bottom dish filled with PHEM buffer. Each measurement was done by acquiring raw images at a series of incident angles (from 0°to 56°with an increment of 4°). ROIs of focal adhesions were determined by threshold or Otsu-based segmentation. The Z-positions were processed and computed using IDL-based custom-written software as described^78^. The topographic images were generated using color code to present the Z-position. The median value of the Z-position of each focal adhesion protein was used as the representative protein position in Z-axis.

### Immunoblotting

Cells were washed with ice-cold PBS and lysed by ice-cold radioimmunoprecipitation assay (RIPA) buffer containing 50mM tris (pH 7.3), 150 mM sodium chloride, 0.25 mM EDTA, 1% Triton X-100, 1% (w/v) sodium deoxycholate, and protease inhibitors cocktail. The lysed cells were incubated on ice for 30min before centrifuge. The protein concentration of the supernatant from each sample was then determined using the Pierce BCA protein assay kit (Thermo Scientific, Catalogue #23225) according to the manufacturer’s instructions. The supernatant with the same amount of protein was mixed with 4X laemmli loading buffer and boiled at 95 – 100℃ for 5 min to denature proteins prior to gel loading. Samples were resolved in 4-20% SurePAGE Bis-Tris gels (GeneScript) and transferred to polyvinylidene difluoride membranes. Membranes were blocked with 5% BSA in Tris-buffered saline with 0.01% Tween 20 (TBS-T) for 1 hour at room temperature and probed with targeted primary antibodies diluted in 5% BSA in TBS-T or PBS-T at 4℃ for overnight. Membranes were then washed for 5 min in TBS-T three times and incubated with HRP-conjugated secondary antibodies for 1 hour at room temperature. Following three times of 5 min TBS-T washes, the Clarity Western ECL substrate (Bio-Rad) was used for developing blots, and the specific proteins were detected and visualized by ChemiDoc Touch (Bio-Rad).

### Co-immunoprecipitation

HEK293T cells were used as the main tool to study the possible mechanism. HEK293T cells were transfected with plasmids tagged with FLAG, HA, or GFP, where anti-FLAG M2 agarose beads (Sigma-Aldrich), anti-HA magnetic beads (ThermoFisher Scientific), or anti-GFP-trap magnetic agarose beads (ChromoTek) were used for each immunoprecipitation according to manufacturer’s instructions. After 24 hours of transfection, cells were lysed using ice-cold RIPA buffer and incubated on ice for 30 min prior to centrifuge. The supernatant was collected and then incubated with equal amount of magnetic beads for immunoprecipitation at 4℃. The magnetic beads were washed with lysis buffer three times and resuspended in 2X laemmli loading buffer, and samples were denatured at 95 – 100℃ for 5 min. Western blotting was conducted for analyzing immunoprecipitation.

### RT-PCR analysis

Qiagen RNeasy mini kit was conducted for harvesting the total RNA of each sample, and cDNA was generated using ThermoFisher Scientific SuperScript IV VILO MasterMix according to manufacturer’s instructions. The equal amount of cDNA was analysed by RT-PCR using SsoFast EvaGreen Supermix (BioRad), and the BioRad CFX96 Touch Real Time PCR machine was performed for measuring the level of cDNA for each given sample. Relative RNA levels were normalized to GAPDH mRNA levels and calculated from ΔCt values compared to its respective control sample. The specific primers used for RT-PCR are listed below:

*BNIP2*: forward 5’- CTGCTTTTTGCGACCTGGC -3’;

*BNIP2*: reverse 5’- AATCCAGGGAGCCAATGTCC -3’;

*TNNT2*: forward 5’- GGAAGAGGCAGACTGAGCGGGA -3’;

*TNNT2*: reverse 5’- TCCCGCGGGTCTTGGAGACTT -3’;

*PTK2*: forward 5’- GCCTTATGACGAAATGCTGGGC-3’;

*PTK2*: reverse 5’- CCTGTCTTCTGGACTCCATCCT -3’;

*PXN*: forward 5’- CTGATGGCTTCGCTGTCGGATT -3’;

*PXN*: reverse 5’- GCTTGTTCAGGTCAGACTGCAG -3’;

*VCL*: forward 5’- TGAGCAAGCACAGCGGTGGATT -3’;

*VCL*: reverse 5’- TCGGTCACACTTGGCGAGAAGA -3’;

*GAPDH*: forward 5’-CCCTTCATTGACCTCAACTACA -3’;

*GAPDH*: reverse 5’-ATGCCAAAGTTGTCATGGAT -3’.

### Plasmid and siRNA transfections

Cells were transfected with jetPRIME (Polyplus-transfection) according to the manufacturer’s instructions. FLAG-, HA-, and GFP-tagged pXJ40 vectors were gifts from E. Manser, Institute for Molecular and Cell Biology, Singapore. Full length human BNIP-2 encoded into pXJ40 vectors were used as templates for generating BNIP-2 truncation constructs, including BNIP-2-cBCH (126-314aa) and BNIP-2-N terminal(1- 166aa). The paxillin, FAK, and vnculin plasmids were from the Davison collection (Kanchanawong laboratory, MBI). High-fidelity DNA polymerase PfuUltra II (Stratagene) was performed for site-directed mutagenesis. Constructs were sequenced to confirm sequence fidelity. The siRNA sequences targeting rat BNIP-2 were purchased from Dharmacon (siBNIP-2-1: 5’-CGUUAGAAGUUAAUGGAAAUU-3’; siBNIP-2-2: 5’-GGAUGAAGGUGGAGAAGUUUU-3’). The RNA interference efficiency was verified using Western blotting to determine the endogenous protein level. Both sequences have 70 to 80% suppression efficiency on BNIP-2. siBNIP-2-1 was used for subsequent experimentation.

### Antibodies

Polyclonal anti-BNIP-2 antibodies were purchased from GeneTex (GTX114283) and Sigma-Aldrich (HPA026843). Anti-glyceraldehyde-3-phosphate dehydrogenase (GAPDH) (sc-47724), monoclonal anti-cardiac troponin T (cTnT) (sc-20025), and mouse IgG (sc-2025) antibodies were purchased from Santa Cruz Biotechnology. Monoclonal anti-myosin light chain 2v (#12975), polyclonal anti-myosin light chain (#3672), polyclonal anti-phospho-myosin light chain 2 (Thr18/Ser19) (#3674), polyclonal anti-src (#2108), polyclonal anti-phospho-src (Tyr416) (#2101), polyclonal anti-FAK (#3285), polyclonal anti-phospho-FAK (Tyr397) (#3283), polyclonal anti-phospho-FAK (Tyr925) (#3284), polyclonal anti-paxillin (#2542), and polyclonal anti-phospho-paxillin (Tyr118) (#2541) antibodies were purchased from Cell Signaling Technology. Monoclonal anti-FAK (#610088) and monoclonal anti-paxillin (#610052) antibodies were from BD Bioscience. Polyclonal anti-vinculin (V4139), polyclonal anti-phospho-vinculin (Tyr822) (V4889), and monoclonal anti-vinculin (V9131) antibodies were from Sigma-Aldrich. Polyclonal anti-GFP (A-11122) antibody was from Life Technology (Invitrogen). The horseradish peroxidase (HRP) secondary antibodies polyclonal antibody against FLAG and polyclonal antibody against HA were from Sigma-Aldrich. All secondary antibodies conjugated with Alexa Fluor dyes for immunostaining were from Life Technologies.

### Proximity ligation assay (PLA)

The proximity ligation assay (Sigma-Aldrich) was performed to detect protein-protein interactions according to manufacturer’s instructions. Briefly, H9c2 cells were seeded on 12 mm glass-bottom dishes overnight and fixed using pre-warmed 4% paraformaldehyde for 15 min at 37℃. After washing in PBS three times, samples were blocked in Duolink blocking solution at 37℃ for 1 hour and incubated with primary antibodies diluted in Duolink antibody diluent at room temperature for 1 hour. After rinsing samples with 1X wash buffer A (Duolink), samples were then incubated with PLUS and MINUS PLA probes diluted in antibody diluent at 37℃ for 1 hour, followed by washing with wash buffer A. DNA Probes were ligated. This circular DNA was amplified and visualized by complementary oligonucleotide probes with a fluorescent label. Samples were mounted with Duolink mounting media with DAPI, and images were acquired using confocal microscopy (60X objective).

### Image analysis

Quantification of immunofluorescence images was performed using ImageJ and MATLAB. Analysis of Western blotting was conducted using ImageLab (Bio-Rad). The area of segmented focal adhesions was measured using Analyze Particles in ImageJ.

### High-throughput RNA sequencing analyses

Total RNA was harvested using Qiagen RNeasy mini kit and sent to BGI Tech Solutions (Hong Kong) Co., Ltd.. Samples were sequenced using DNBseq platform with 30 million sequencing depth. Differential expression was calculated with DESeq2. BGI Dr. Tom analysis platform was conducted for generating clustered heatmap and weighted correlation network analysis. PCA scores plot was generated using the principal components analysis in R studio. FPKM was performed for all measurements.

### Statistical analysis

The Western blotting and immunofluorescence data were presented as means ± SEM. The statistic graph of SAIM results was generated using OriginPro 2019. The interquartile range was presented for each focal adhesion protein, and the median value was labeled in the graph. All statistical tests were created using GraphPad Prism 9.1.1. Differences were considered statistically significant if the P-value is less than 0.05, where \**P*<0.05, \*\**P*<0.01, \*\*\**P*<0.001 and \*\*\*\**P*<0.0001. At least three independent biological repeats were performed for all experiments involving statistical comparison.

## Data availability

All study data are included in the article and/or SI appendix. Working models were created with BioRender.com.

## ACKNOWLEDGEMENTS

We thank core facilities of Mechanobiology Institute of Singapore for the support and assistance provided.

## AUTHOR CONTRIBUTIONS

J.X., J.W.A., D.C.P.W., X.Z., C.J.M.L., P.K., and B.C.L. conceived and designed the experiments. J.X., J.W.A., D.C.P.W., X.Z., and C.J.M.L. conducted the experiments. X.Z. performed and analyzed the scanning angle interference microscopy experiments. C.J.M.L. performed the hESC cardiomyocyte differentiation and CRISPR-targeted knockdown. J.X., J.W.A., P.K., and B.C.L. wrote the manuscript. All authors discussed the results and commented on it.

## Supporting Information for

**Fig. S1.**
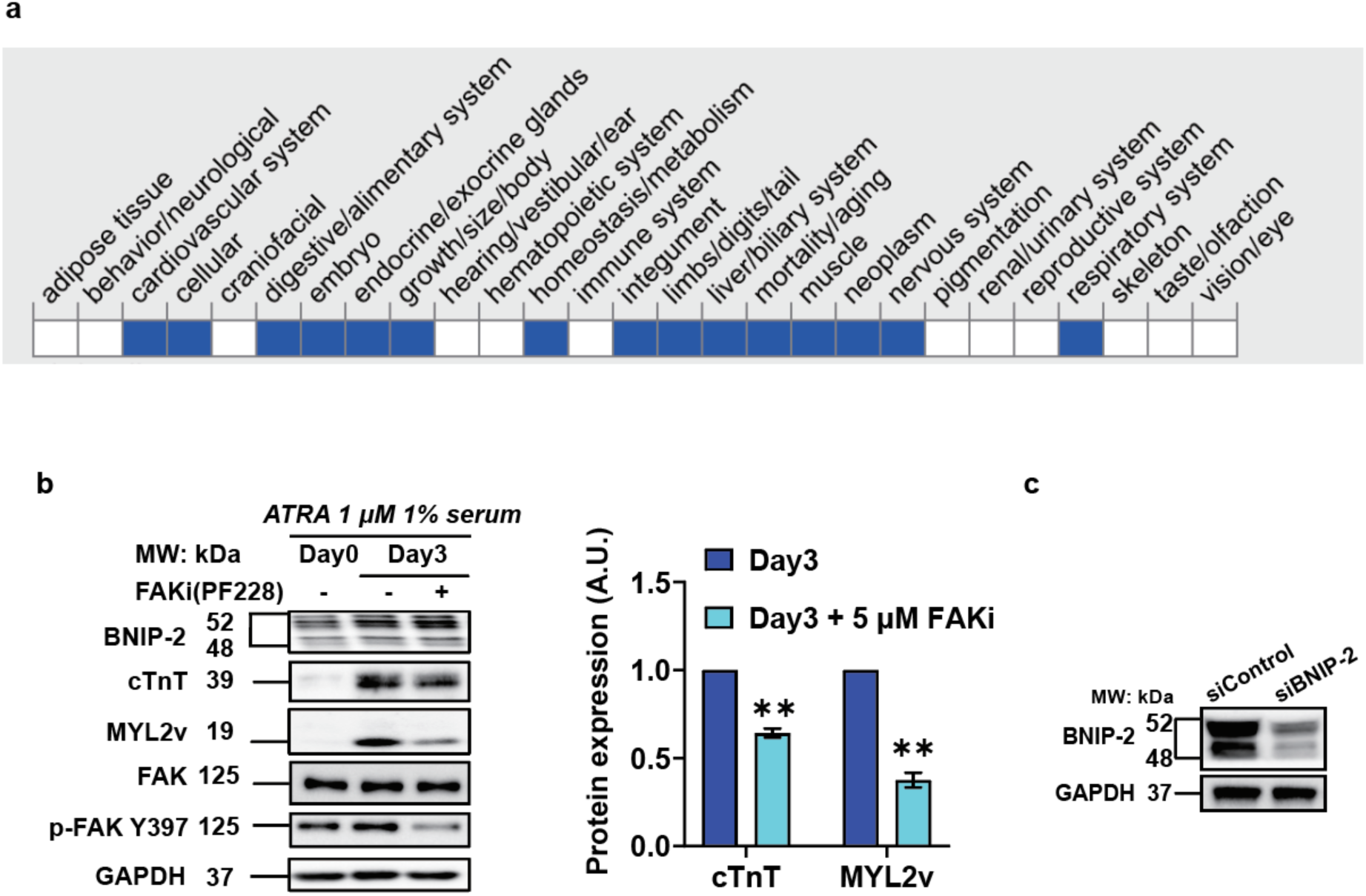
FAK inhibition impedes cardiac differentiation. **a**, The lack of FAK in mice results in the dysfunction of cardiovascular system, among others. Data retrieved from Mouse Genome Database (MGD)(1). **b**, Immunoblot analyses of BNIP-2, cardiac troponin T (cTnT), myosin light chain 2v (MYL2v), FAK, phosphorylated FAK Y397 in differentiated H9c2 cells (Left). Cells were treated with 5 μM FAK inhibitor and 1 μM ATRA in low serum medium (1% FBS) for three days. Level of cTnT and MYL2v relative to Day 3 cells under control condition (right). Mean ± s.e.m., *n*=3 independent experiments. Unpaired two-tailed Student’s t-test; \**P*<0.05, \*\**P*<0.01, and \*\*\**P*<0.001. **c**, Verification of BNIP-2 knockdown efficiency in H9c2 cells 48 hours after siRNA transfection.

**Fig. S2.**
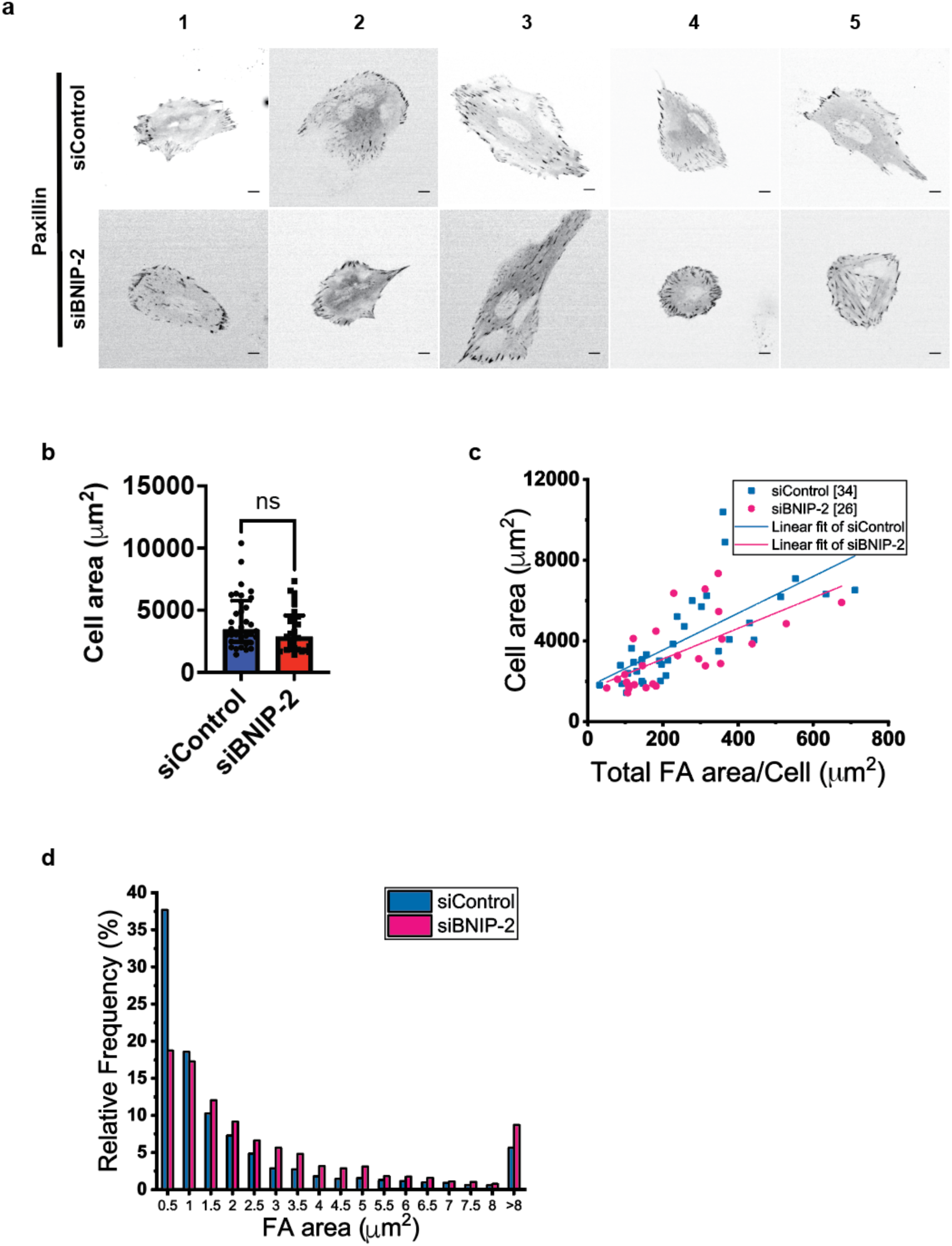
BNIP-2 reduction increases FA area in H9c2 myoblasts. **a**, Immunofluorescence analysis of FA area in BNIP-2 knockdown H9c2 cells. Scale bar, 10 μm. **b**, Quantification of cell area in control and BNIP-2 knockdown cells. Each data point represents a single cell. *n* = 34 (siControl) and 26 (siBNIP-2) cells. Median ± interquartile. Unpaired two-tailed Student’s t-test; **c**, Polynomial linear fitting curve of cell area and total area of FAs per cell in siControl (blue curve) and siBNIP-2 (red curve) H9c2 cells. Each data point presents individual cells in four independent experiments. **d**, Histogram of FA sizes in control and BNIP-2 knockdown cells.

**Fig. S3.**
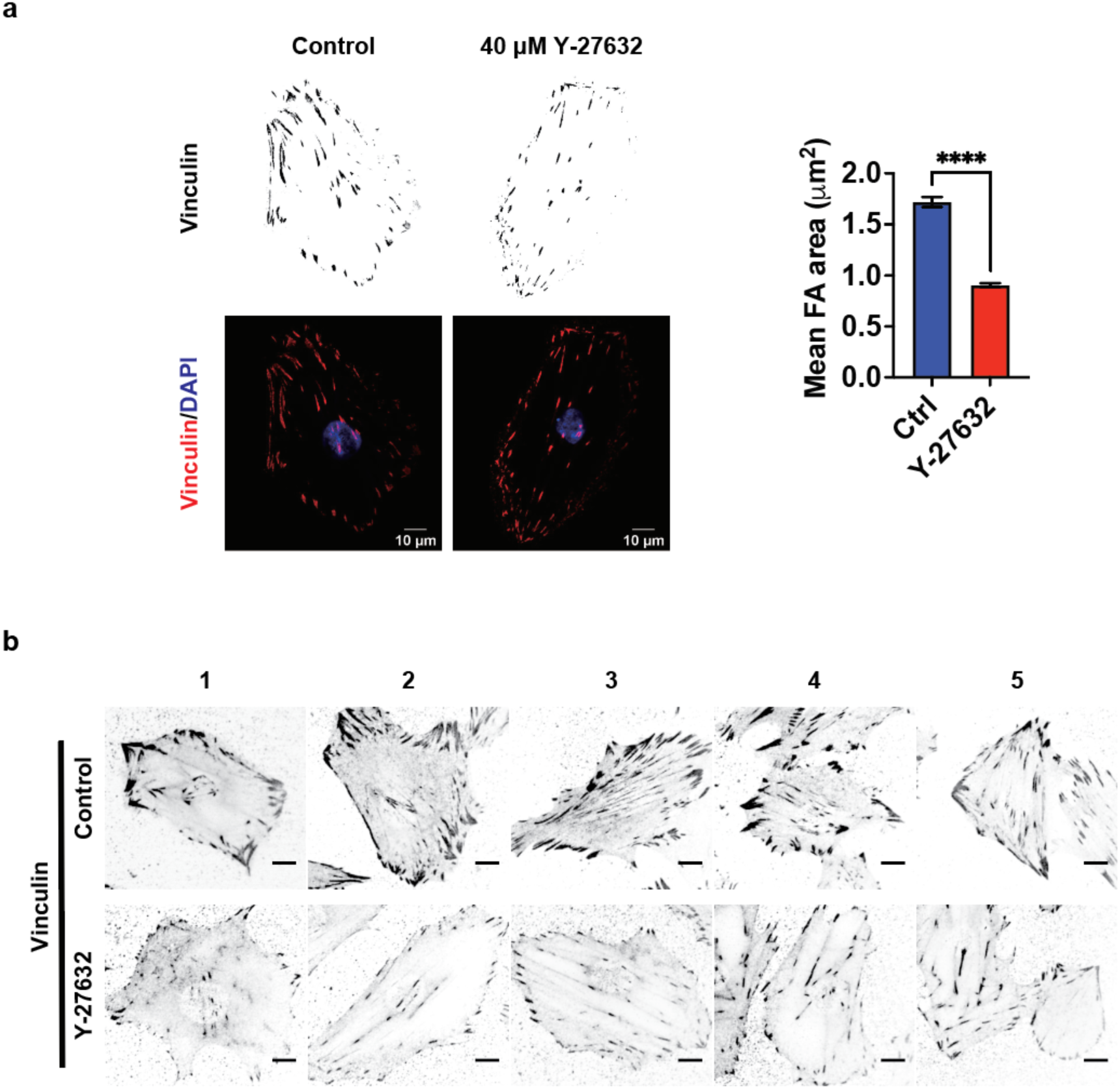
ROCK inhibition reduces FA size in H9c2 myoblasts. **a**, Immunofluorescence analysis of FA area in H9c2 myoblasts treated with ROCK inhibitor (Y-27632) (left). Scale bar, 10 μm. Quantification of FA area is shown (right). Each data point represents every FA in cells. Mean ± s.e.m., *n* = 3308 FAs from 20 cells (control) and 4928 FAs from 23 cells (Y-27632). Unpaired two-tailed Student’s t-test; \*\*\*\**P*<0.0001. **b**, Immunofluorescence analysis of FA area in H9c2 cells treated with Y-27632. Scale bar, 10 μm.

**Fig. S4.**
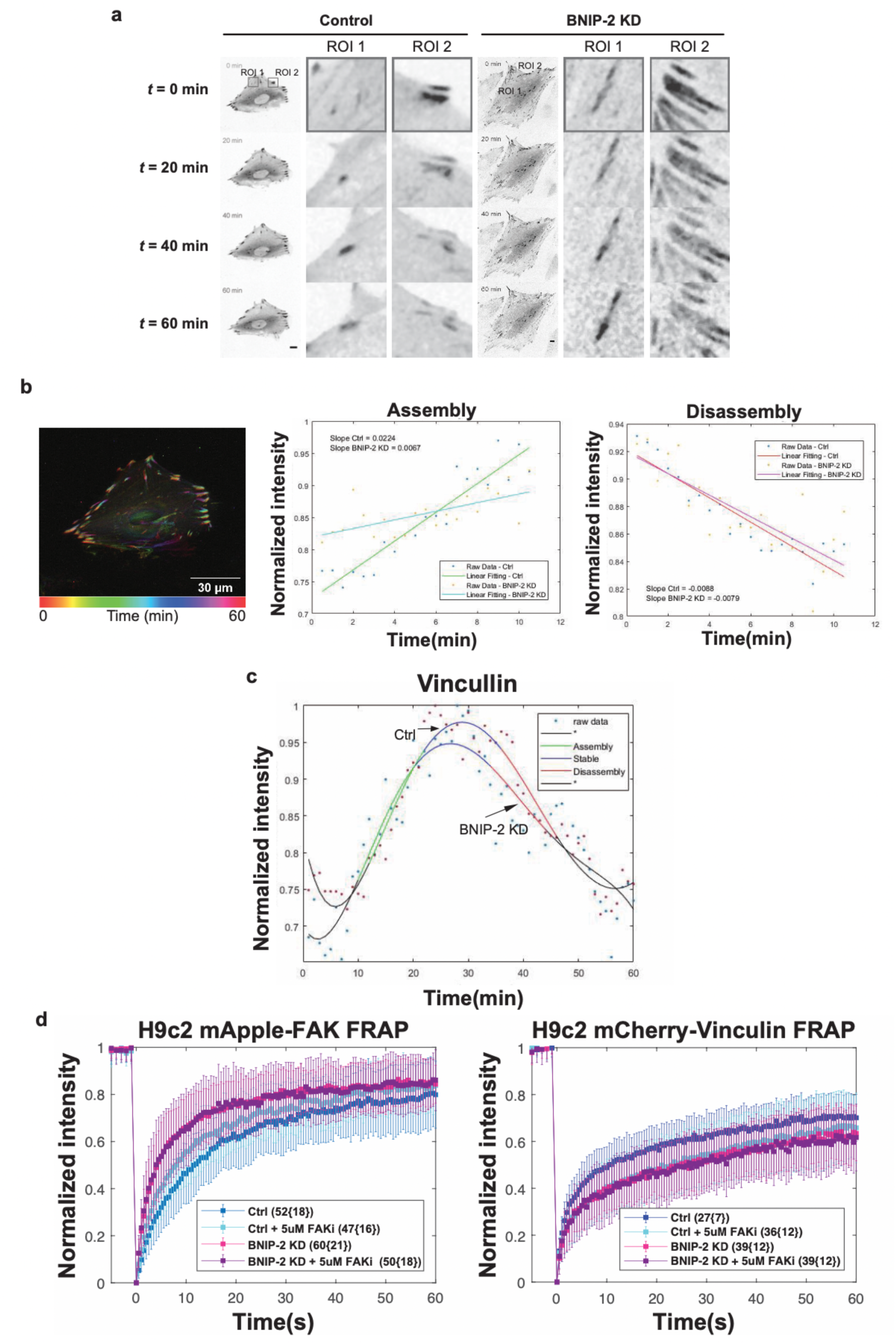
BNIP-2 controls FA dynamics in H9c2 myoblasts. **a**, Time-lapse images of control and BNIP-2 knockdown H9c2 cells expressing mApple-paxillin. Scale bar, 10 μm. **b**, The overlaying time-lapse images of H9c2 cells expressing mApple-paxillin. Each time point is labeled with a different color. Scale bar, 30 μm (left). Calculation of assembly (middle) and disassembly (right) rates fitted with polynomial linear model. **c,** Time-lapse imaging for measuring assembly and disassembly rates of FA proteins in H9c2 cells expressing mCherry-vinculin and siRNA-targeting BNIP-2. **d,** FRAP-relative fluorescence intensity curves of control, BNIP-2-knockdown, and FAK-inhibited H9c2 cells expressing mApple-FAK (left) and mCherry-vinculin (right).

**Fig. S5.**
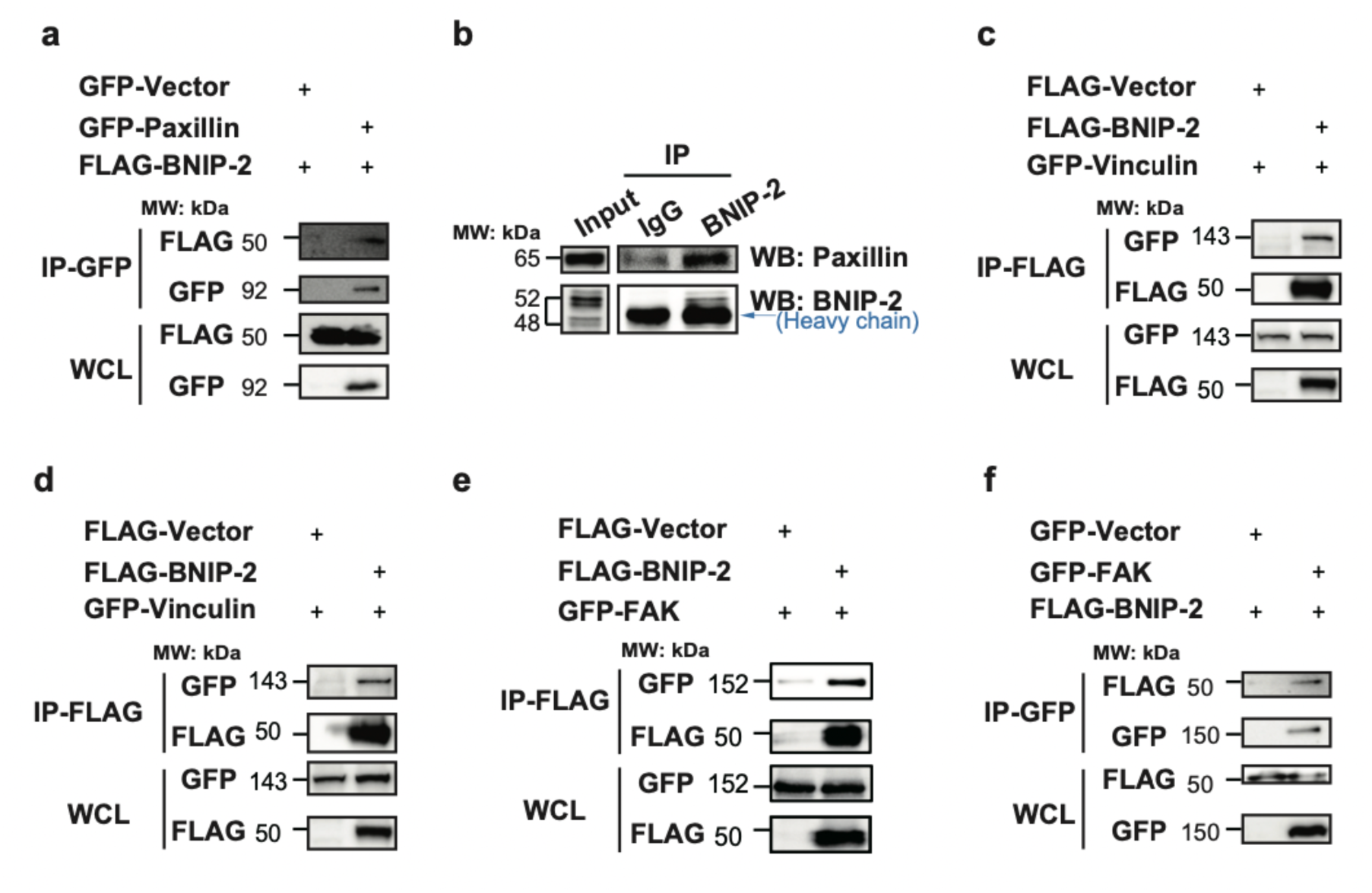
Co-IP replicates for determining interactions between BNIP-2 and FA proteins. **a**, Detection of interactions between BNIP-2 and paxillin in HEK293T cells expressing GFP-tagged empty vector, paxillin, and FLAG-BNIP-2. Anti-GFP-trap magnetic agarose beads were used for immunoprecipitation according to manufacturer’s instruction. The immunoprecipitation blot detected using anti-FLAG antibody indicates the interaction between BNIP-2 and paxillin. **b**, BNIP-2 forms a physiologic complex with paxillin in H9c2 cells. The cell lysates were incubated with BNIP-2 antibody or immunoglobulin G (IgG) control for endogenous immunoprecipitation using Protein A/G PLUS agarose beads. Western blotting was performed with paxillin and BNIP-2 antibodies. The blue arrow denotes heavy chains. The immunoprecipitation blot detected using anti-paxillin antibody indicates the interaction between endogenous BNIP-2 and paxillin. **c**-**d**, Detection of interactions between BNIP-2 and vinculin in HEK293T cells expressing FLAG-tagged empty vector, BNIP-2, and GFP-vinculin. Anti-FLAG magnetic agarose beads were used for immunoprecipitation according to manufacturer’s instruction. The immunoprecipitation blot detected using anti-GFP antibody indicates the interaction between BNIP-2 and vinculin. **e**-**f**, Detection of interaction between BNIP-2 and FAK in HEK293T cells expressing FLAG-tagged empty vector, BNIP-2, and GFP-FAK (**e**), or GFP-tagged empty vector, FAK, and FLAG-BNIP-2 (**f**). Anti-FLAG (**e**) or anti-GFP-trap (**f**) magnetic agarose beads were used for immunoprecipitation according to manufacturer’s instruction. The immunoprecipitation blot detected using anti-GFP (**e**) or anti-FLAG (**f**) antibody indicates the interaction between BNIP-2 and FAK.

**Fig. S6.**
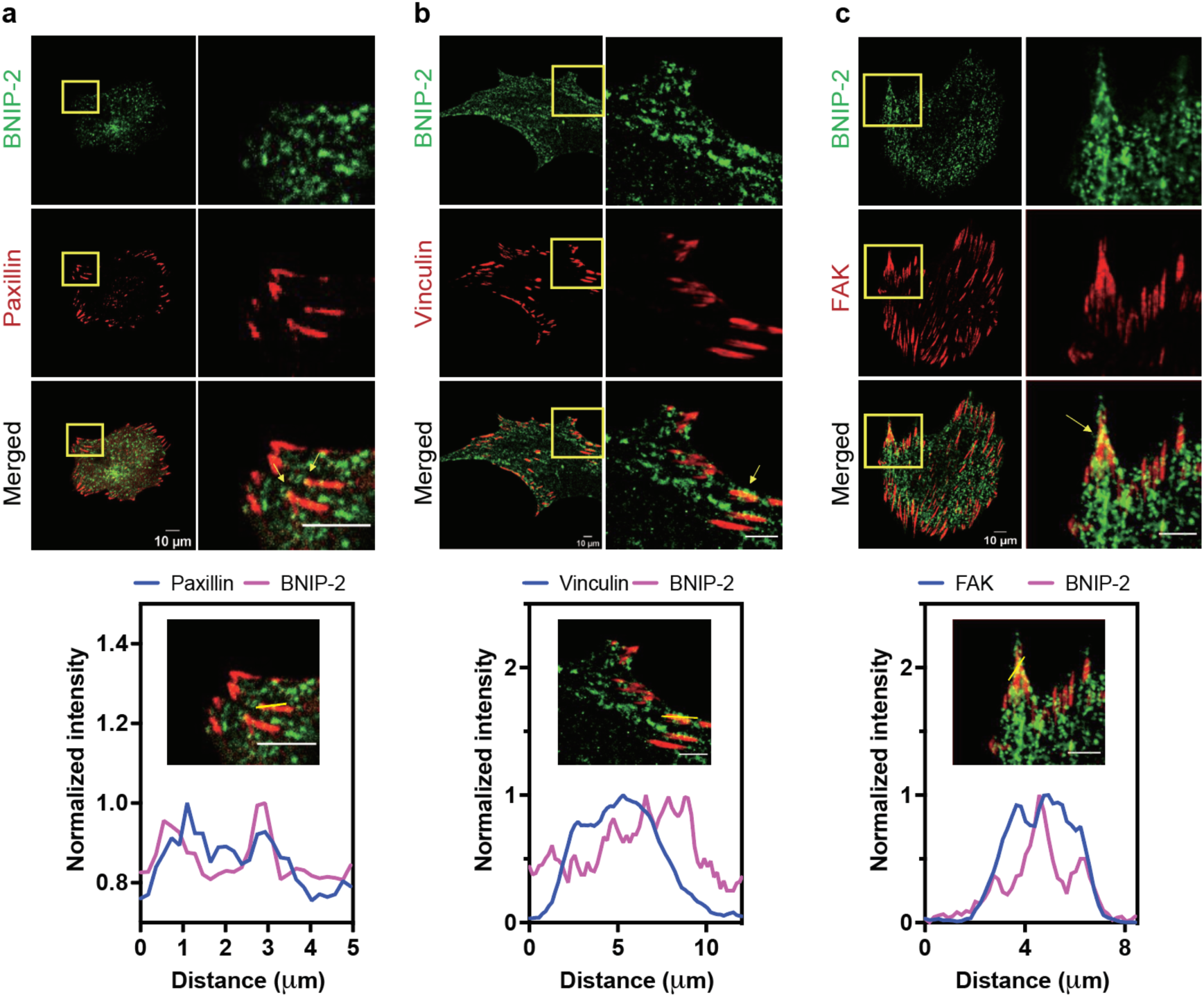
BNIP-2 colocalized with paxillin, vinculin, and FAK. **a-c**, Images showing the localization of paxillin, vinculin, and FAK (from left to right) together with the staining of BNIP-2 in H9c2 myoblasts. Zoomed ROIs are shown (right). Histograms (bottom) show the normalized fluorescence intensity of BNIP-2 and FA proteins along the yellow line in the merged images. Arbitrary units. Scale bar, 10 μm.

**Fig. S7.**
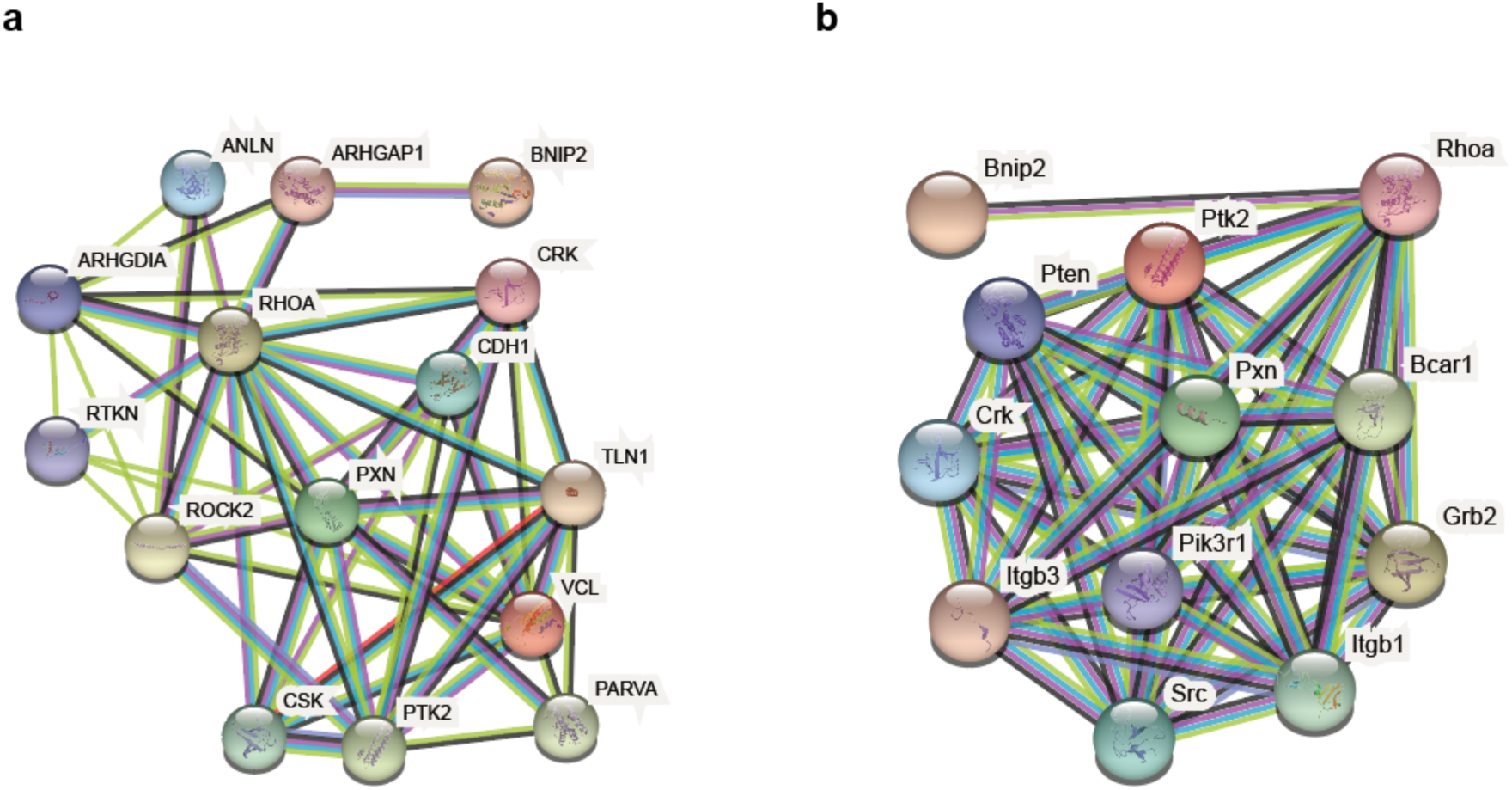
Interaction networks of BNIP-2 and FA proteins in human and rats. **a-b**, Protein-protein interactions between BNIP-2 and FA proteins in human (**a**) and rat (**b**) retrieved from STRING(https://string-db.org/). PXN/Pxn: Paxillin; PTK2/Ptk2: FAK; VCL/Vcl: Vinculin; BNIP2/Bnip2: BNIP-2.

**Fig. S8.**
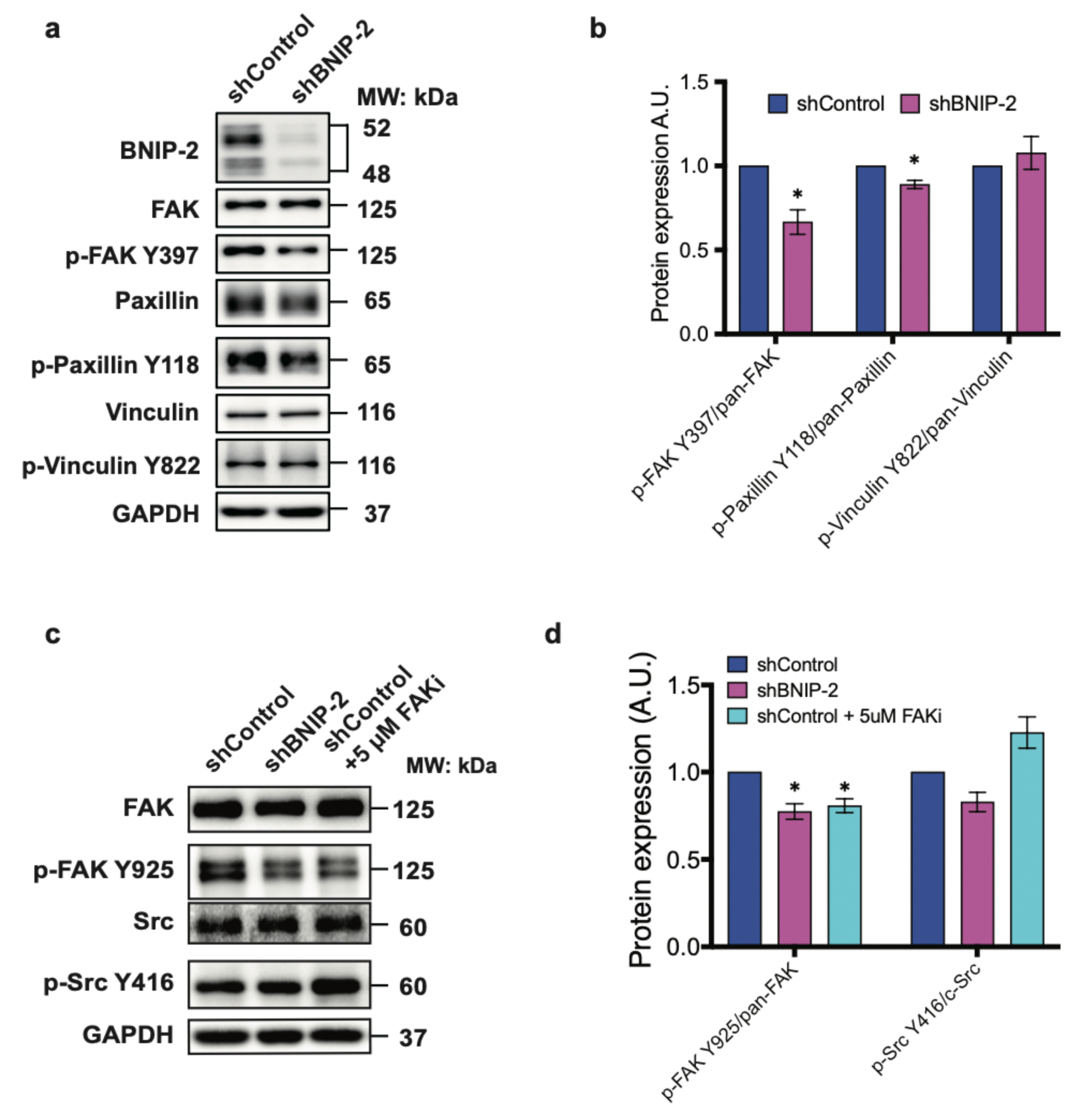
BNIP-2 knockdown reduces the phosphorylation of FAK and paxillin. **a-b**, Immunoblot analyses of BNIP-2, FAK, paxillin, vinculin, phosphorylated FAK Y397, p-paxillin Y118, and p-vinculin Y822 in control and BNIP-2 knockdown H9c2 cells. The phosphorylated level was normalized to the total protein expression of each given protein. **c**-**d**, Immunoblot analyses of FAK, src, phosphorylated FAK Y925, and p-Src Y416 in H9c2 cells under conditions of control, BNIP-2 knockdown, and FAK inhibition. The phosphorylated level was normalized to the total protein expression of each given protein. Mean ± s.e.m., *n*=3 independent experiments. Unpaired two-tailed Student’s t-test; \**P*<0.05.

**Fig. S9.**
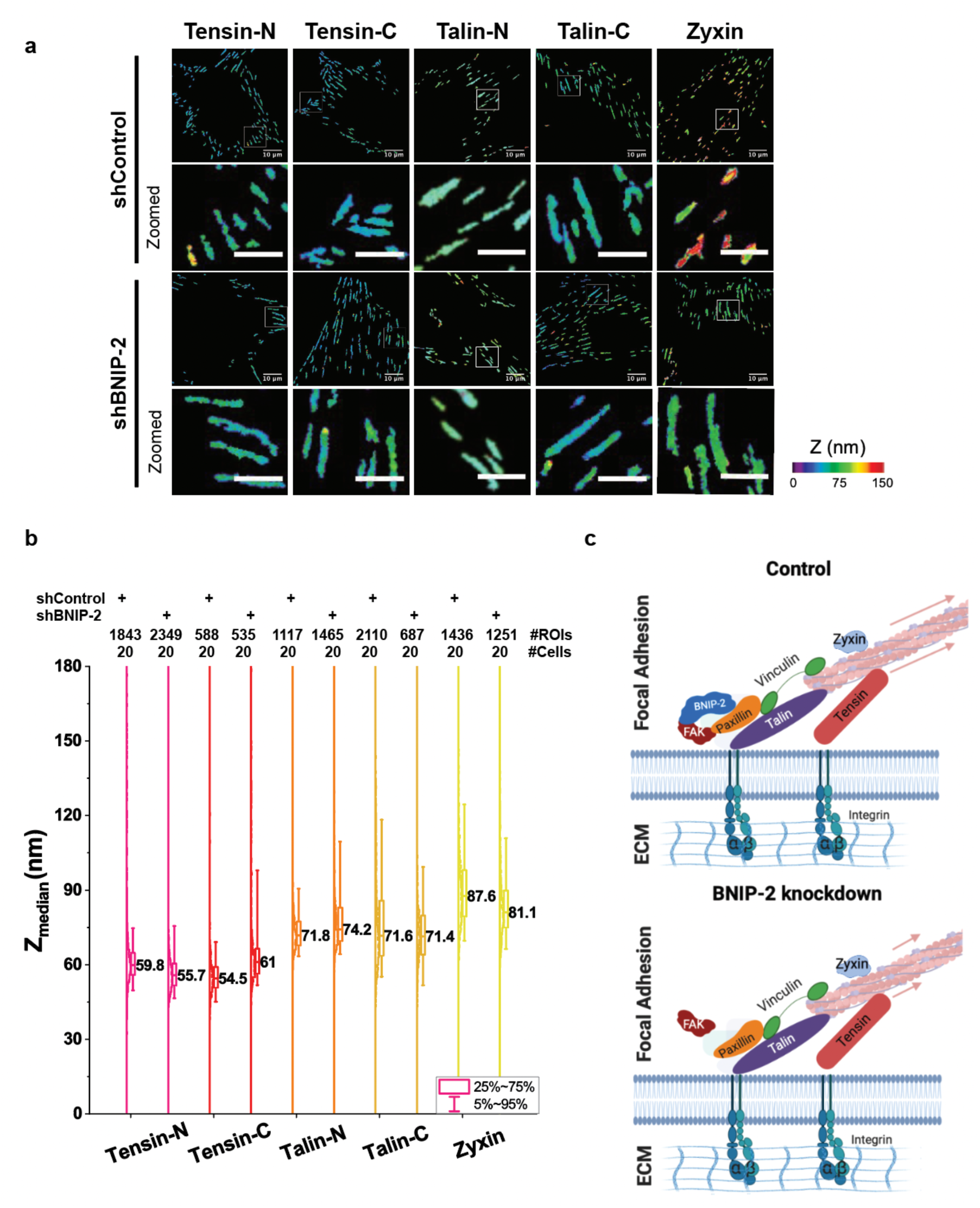
The BNIP-2 regulation of FA nanostructure. **a**, Color-coded topographic map in Z-axis of FA proteins in control and BNIP-2 knockdown H9c2 cells expressing fluorophore-tagged tensin-N, tensin-C, talin-N, talin-C, and zyxin. Color bar indicates the Z-position relative to the surface. Scale bar, 10 μm (original) and 5 μm (zoomed). **b**, Histogram and box plot of FA Z-positions. Each data point represents the Z-center of individual FA region of interest (ROI) relative to the surface. The number of ROIs and cells labelled at the top of the plot. Median ± interquartile. **c**, Schematic diagram of FA nanostructure in control (top) and BNIP-2 knockdown (bottom) H9c2 myoblasts.

**Fig. S10.**
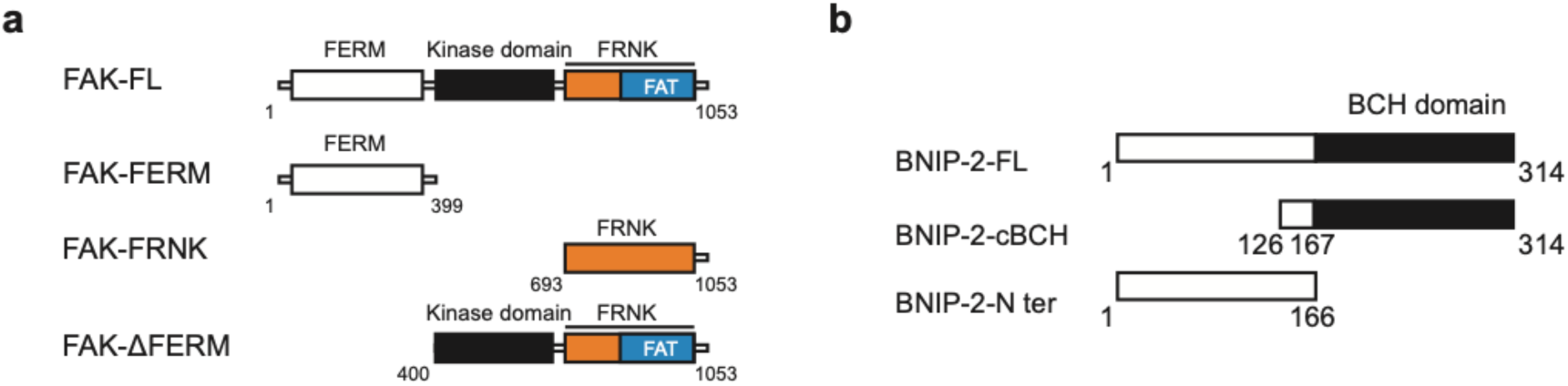
The schematic diagrams of truncation mutants of FAK and BNIP-2. **a**, Region illustrations of FAK truncated mutants (Gallus gallus). FAK-FL: FAK full-length; FAK-FERM: FAK FERM domain with the linker between FERM and kinase domain; FAK-FRNK: FAK FRNK domain. **b**, Region illustrations of BNIP-2 truncated mutants (Homo Sapiens). BNIP-2-FL: BNIP-2 full-length; BNIP-2-cBCH: BNIP-2 cBCH domain; BNIP-2-N ter: BNIP-2 N-terminal domain.

**Fig. S11.**
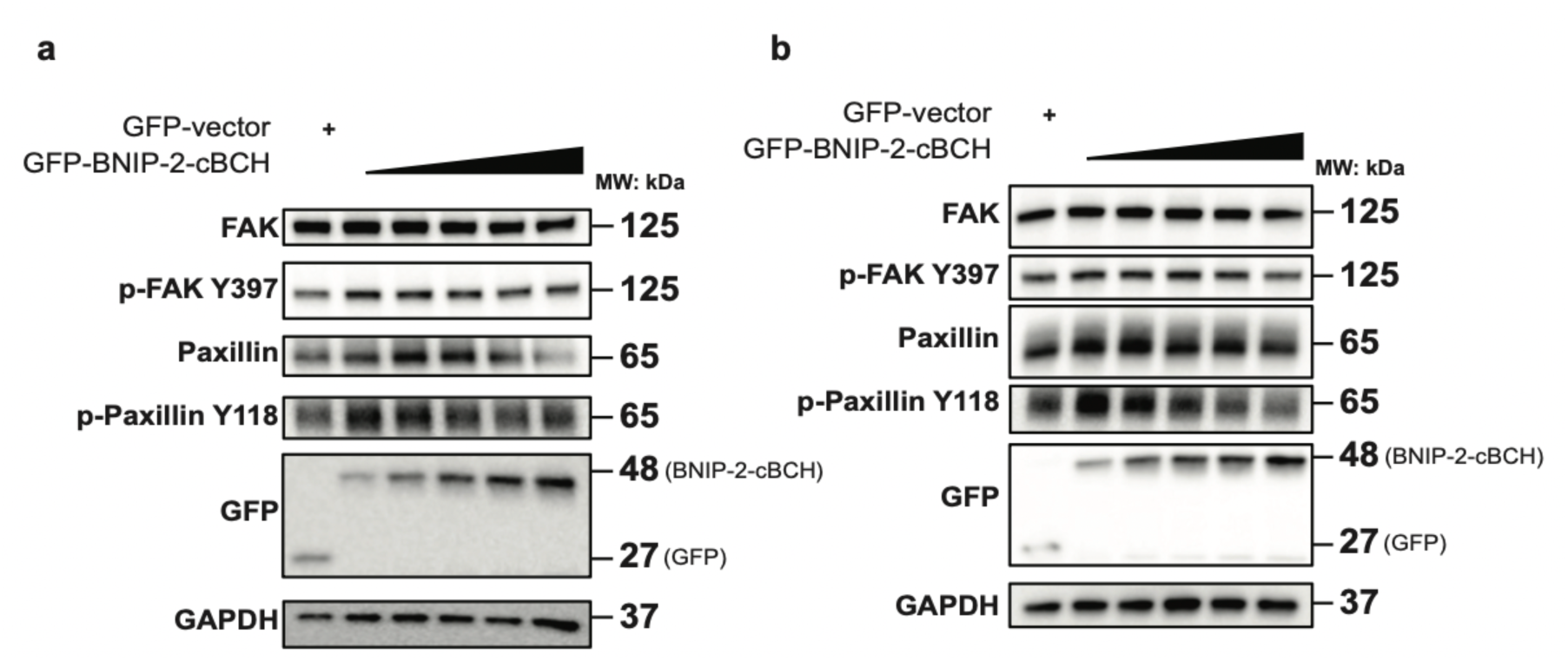
BNIP-2 has a scaffolding effect for FAK/paxillin signalling. **a-b**, Immunoblot analyses of FAK, phosphorylated FAK Y397, paxillin, phosphorylated paxillin Y118, and GFP in H9c2 myoblasts expressing gradient amount of GFP-BNIP-2-cBCH (illustrated by black triangle). Anti-GFP blot denotes the expression of GFP-BNIP-2-cBCH.

**Fig. S12.**
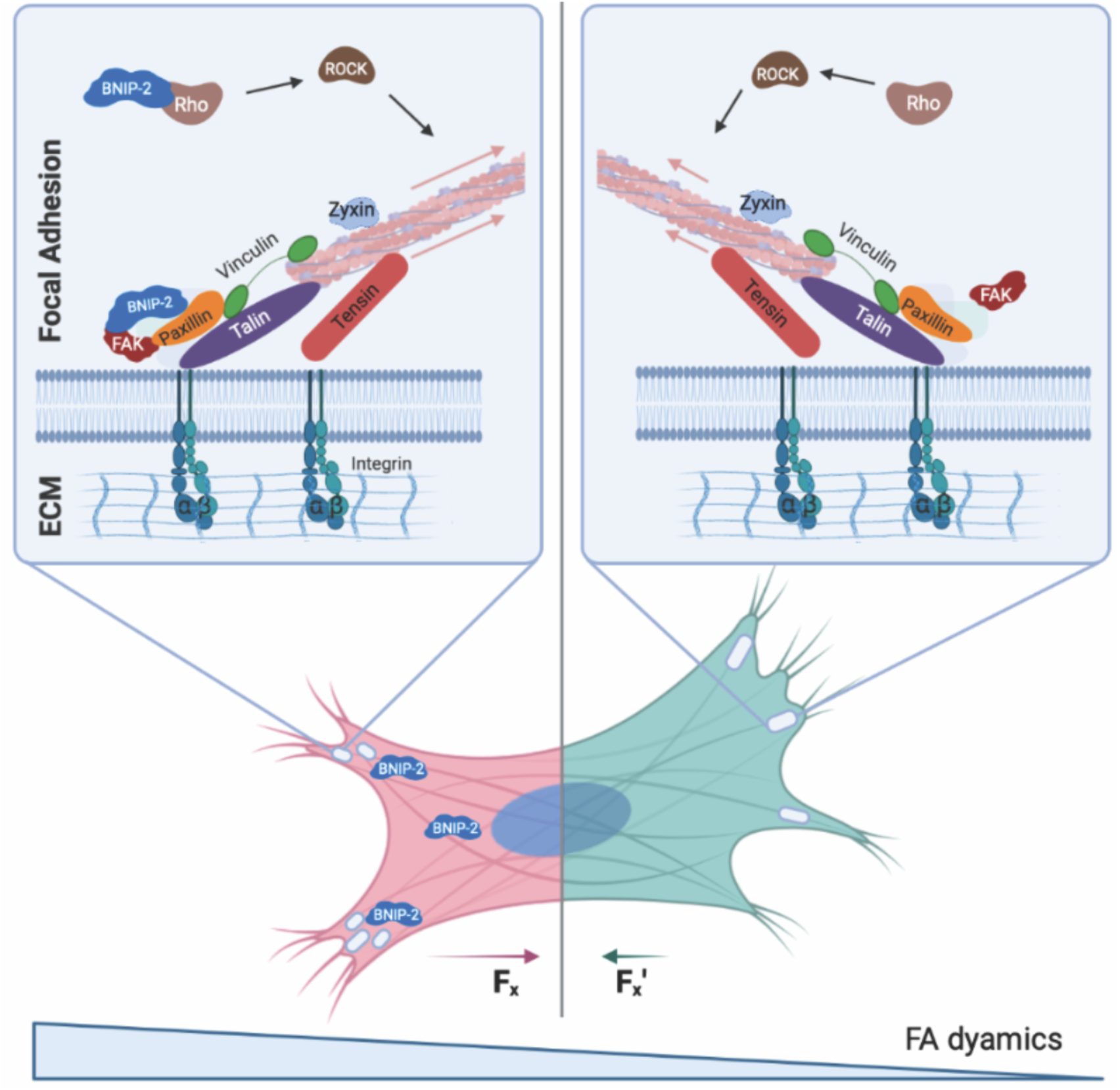
Summary model of BNIP-2 action at FAs that directs FAK local organization and FA signaling. BNIP-2 interacts with FAK through its BCH domain to regulate FAK/paxillin signaling and facilitate mechanotransduction. BNIP-2 acts as a scaffold for FAK/paxillin and paxillin/vinculin interactions. The reduction of BNIP-2 induces larger FAs, reduced FA dynamics, and lower contractility in H9c2 cells. We propose that BNIP-2 coordinates the integration of FAK, paxillin and vinculin at FAs and the local spatial distribution of FAK, leading to the activation of FAK on paxillin signaling to ensure downstream force transmission, thus supporting H9c2 and hESC cardiomyocyte differentiation.

## References

1. Granados-Riveron, J.T. & Brook, J.D. The Impact of Mechanical Forces in Heart Morphogenesis. Circulation: Cardiovascular Genetics 5, 132–142 (2012).

2. Hove, J.R. et al. Intracardiac fluid forces are an essential epigenetic factor for embryonic cardiogenesis. Nature (2003).

3. Smith, L.R., Cho, S. & Discher, D.E. Stem Cell Differentiation is Regulated by Extracellular Matrix Mechanics. Physiology (Bethesda) 33, 16–25 (2018).

4. Midgett, M. & Rugonyi, S. Congenital heart malformations induced by hemodynamic altering surgical interventions. Frontiers in Physiology (2014).

5. Kanchanawong, P. & Calderwood, D.A. Organization, dynamics and mechanoregulation of integrin-mediated cell–ECM adhesions. Nature Reviews Molecular Cell Biology (2022).

6. Argentati, C. et al. Insight into Mechanobiology: How Stem Cells Feel Mechanical Forces and Orchestrate Biological Functions. International Journal of Molecular Sciences 20 (2019).

7. Romaine, A. et al. Overexpression of integrin α11 induces cardiac fibrosis in mice. Acta Physiologica 222, e12932 (2018).

8. Higashikuse, Y. et al. Perturbation of the titin/MURF1 signaling complex is associated with hypertrophic cardiomyopathy in a fish model and in human patients. Disease Models & Mechanisms 12, dmm041103 (2019).

9. Münch, J. & Abdelilah-Seyfried, S. Sensing and Responding of Cardiomyocytes to Changes of Tissue Stiffness in the Diseased Heart. Frontiers in Cell and Developmental Biology 9 (2021).

10. Lucitti, J.L. et al. Vascular remodeling of the mouse yolk sac requires hemodynamic force. Development (2007).

11. Andrés-Delgado, L. & Mercader, N. Interplay between cardiac function and heart development. Biochimica et Biophysica Acta (BBA) - Molecular Cell Research 1863, 1707–1716 (2016).

12. Kanchanawong, P. et al. Nanoscale architecture of integrin-based cell adhesions. Nature 468, 580–584 (2010).

13. Sun, Z.A.-O., Guo, S.S. & Fässler, R. Integrin-mediated mechanotransduction. Journal of cell biology (2016).

14. Case, L.B. & Waterman, C.M. Integration of actin dynamics and cell adhesion by a three-dimensional, mechanosensitive molecular clutch. Nature Cell Biology 17, 955–963 (2015).

15. Crocini, C. & Gotthardt, M. Cardiac sarcomere mechanics in health and disease. Biophysical Reviews 13, 637–652 (2021).

16. Adamo, L. et al. Biomechanical forces promote embryonic haematopoiesis. Nature 459, 1131–1135 (2009).

17. Samarel, A.M. Costameres, focal adhesions, and cardiomyocyte mechanotransduction. American Journal of Physiology-Heart and Circulatory Physiology 289, H2291–H2301 (2005).

18. Yim, E.K.F. & Sheetz, M.P. Force-dependent cell signaling in stem cell differentiation. Stem Cell Research & Therapy 3, 41 (2012).

19. Pasapera, A.M., Schneider, I.C., Rericha, E., Schlaepfer, D.D. & Waterman, C.M. Myosin II activity regulates vinculin recruitment to focal adhesions through FAK-mediated paxillin phosphorylation. Journal of Cell Biology 188, 877–890 (2010).

20. Subauste, M.C. et al. Vinculin modulation of paxillin-FAK interactions regulates ERK to control survival and motility. Journal of Cell Biology (2004).

21. Clemente, C.F.M.Z. et al. Focal adhesion kinase governs cardiac concentric hypertrophic growth by activating the AKT and mTOR pathways. Journal of molecular and cellular cardiology 52 **2**, 493–501 (2012).

22. Peng, X. et al. Inactivation of focal adhesion kinase in cardiomyocytes promotes eccentric cardiac hypertrophy and fibrosis in mice. The Journal of clinical investigation 116 **1**, 217–227 (2006).

23. Peng, X. et al. Cardiac developmental defects and eccentric right ventricular hypertrophy in cardiomyocyte focal adhesion kinase (FAK) conditional knockout mice. Proceedings of the National Academy of Sciences 105, 6638 (2008).

24. Ohashi, K., Fujiwara, S. & Mizuno, K. Roles of the cytoskeleton, cell adhesion and rho signalling in mechanosensing and mechanotransduction. The Journal of Biochemistry 161, 245–254 (2017).

25. Wolfenson, H., Bershadsky, A.D., Henis, Y.I. & Geiger, B. Actomyosin-generated tension controls the molecular kinetics of focal adhesions. Journal of Cell Science 124, 1425 - >1432 (2011).

26. Dumbauld, D.W. et al. How vinculin regulates force transmission. Proceedings of the National Academy of Sciences 110, 9788 - 9793 (2013).

27. Grashoff, C. et al. Measuring mechanical tension across vinculin reveals regulation of focal adhesion dynamics. Nature 466, 263–266 (2010).

28. Tham, Y.K., Bernardo, B.C., Ooi, J.Y.Y., Weeks, K.L. & McMullen, J.R. Pathophysiology of cardiac hypertrophy and heart failure: signaling pathways and novel therapeutic targets. Archives of Toxicology 89, 1401–1438 (2015).

29. Wong, D.C.P. et al. BNIP-2 Activation of Cellular Contractility Inactivates YAP for H9c2 Cardiomyoblast Differentiation. Advanced Science 9, 2202834 (2022).

30. Pan, M. et al. BNIP-2 retards breast cancer cell migration by coupling microtubule-mediated GEF-H1 and RhoA activation. Science Advances 6, eaaz1534 (2020).

31. Yi, P. et al. KIF5B transports BNIP-2 to regulate p38 mitogen-activated protein kinase activation and myoblast differentiation. Molecular biology of the cell (2015).

32. Pan, C.Q. & Low, B.C. Functional plasticity of the BNIP-2 and Cdc42GAP Homology (BCH) domain in cell signaling and cell dynamics. FEBS Letters 586, 2674–2691 (2012).

33. Chichili Vishnu Priyanka, R., et al. Structural basis for p50RhoGAP BCH domain– mediated regulation of Rho inactivation. Proceedings of the National Academy of Sciences 118, e2014242118 (2021).

34. Majkut, S. et al. Heart-Specific Stiffening in Early Embryos Parallels Matrix and Myosin Expression to Optimize Beating. Current Biology 23, 2434–2439 (2013).

35. Guo, Y. & Pu, W.T. Cardiomyocyte Maturation. Circulation Research 126, 1086–1106 (2020).

36. Bershadsky, A.D., Balaban, N.Q. & Geiger, B. Adhesion-Dependent Cell Mechanosensitivity. Annual Review of Cell and Developmental Biology 19, 677–695 (2003).

37. Bult, C.J. et al. Mouse Genome Database (MGD) 2019. Nucleic Acids Research 47, D801–D806 (2019).

38. Bult, C.J., Blake, J.A., Smith, C.L., Kadin, J.A. & Richardson, J.E. Mouse Genome Database (MGD) 2019.

39. Zordoky, B.N. & El-Kadi, A.O. H9c2 cell line is a valuable in vitro model to study the drug metabolizing enzymes in the heart. Journal of pharmacological and toxicological methods (2007).

40. Branco, A.F. et al. Gene Expression Profiling of H9c2 Myoblast Differentiation towards a Cardiac-Like Phenotype. PLOS ONE (2015).

41. Balaban, N.Q. et al. Force and focal adhesion assembly: a close relationship studied using elastic micropatterned substrates. Nature Cell Biology 3, 466–472 (2001).

42. Coniglio, S., Zavarella, S. & Symons, M. Pak1 and Pak2 Mediate Tumor Cell Invasion through Distinct Signaling Mechanisms. Molecular and cellular biology 28, 4162–4172 (2008).

43. Goetsch, K., Snyman, C., Myburgh, K. & Niesler, C. ROCK-2 is Associated With Focal Adhesion Maturation During Myoblast Migration. Journal of Cellular Biochemistry 115 (2014).

44. De Pascalis, C. & Etienne-Manneville, S. Single and collective cell migration: The mechanics of adhesions. Molecular Biology of the Cell 28, 1833–1846 (2017).

45. Stutchbury, B.A.-O., Atherton, P.A.-O.X., Tsang, R.A.-O., Wang, D.A.-O. & Ballestrem, C.A.-O. Distinct focal adhesion protein modules control different aspects of mechanotransduction. J Cell Sci (2017).

46. Yao, M. et al. Mechanical activation of vinculin binding to talin locks talin in an unfolded conformation. Scientific Reports 4, 4610 (2014).

47. Hamadi, A. et al. Regulation of focal adhesion dynamics and disassembly by phosphorylation of FAK at tyrosine 397. Journal of Cell Science 118, 4415–4425 (2005).

48. Swaminathan, V., Fischer, R.S. & Waterman, C.M. The FAK-Arp2/3 interaction promotes leading edge advance and haptosensing by coupling nascent adhesions to lamellipodia actin. Molecular biology of the cell 27, 1085–1100 (2016).

49. Yamada, K.M. & Geiger, B. Molecular interactions in cell adhesion complexes. Current Opinion in Cell Biology 9, 76–85 (1997).

50. Szklarczyk, D. et al. STRING v11: protein–protein association networks with increased coverage, supporting functional discovery in genome-wide experimental datasets. Nucleic Acids Research 47, D607–D613 (2019).

51. Zaidel-Bar, R., Itzkovitz, S., Ma’ayan, A., Iyengar, R. & Geiger, B. Functional atlas of the integrin adhesome. Nat Cell Biol 9, 858–867 (2007).

52. Turner, C.E. Paxillin and focal adhesion signalling. Nature Cell Biology 2, E231–E236 (2000).

53. Deramaudt, T.B. et al. Altering FAK-paxillin interactions reduces adhesion, migration and invasion processes. PLOS ONE (2014).

54. Blaukat, A. Focal Adhesion Kinase, in xPharm: The Comprehensive Pharmacology Reference. (eds. S.J. Enna & D.B. Bylund) 1–10 (Elsevier, New York; 2007).

55. Mitra, S.K. & Schlaepfer, D.D. Integrin-regulated FAK–Src signaling in normal and cancer cells. Current Opinion in Cell Biology 18, 516–523 (2006).

56. Ziegler, W.H., Liddington Rc Fau - Critchley, D.R. & Critchley, D.R. The structure and regulation of vinculin. Trends in cell biology (2006).

57. López-Colomé, A.M., Lee-Rivera, I., Benavides-Hidalgo, R. & López, E. Paxillin: a crossroad in pathological cell migration. Journal of Hematology & Oncology 10, 50 (2017).

58. Xia, S., Yim, E. & Kanchanawong, P. Molecular Organization of Integrin-Based Adhesion Complexes in Mouse Embryonic Stem Cells. ACS Biomaterials Science & Engineering 5 (2019).

59. Paszek, M.J. et al. Scanning angle interference microscopy reveals cell dynamics at the nanoscale. Nature Methods (2012).

60. Yoon, H., Dehart, J.P., Murphy, J.M. & Lim, S.-T.S. Understanding the Roles of FAK in Cancer: Inhibitors, Genetic Models, and New Insights. Journal of Histochemistry & Cytochemistry 63, 114–128 (2014).

61. Linari, M. et al. Force generation by skeletal muscle is controlled by mechanosensing in myosin filaments. Nature 528, 276–279 (2015).

62. Kampourakis, T., Sun, Y.-B. & Irving, M. Myosin light chain phosphorylation enhances contraction of heart muscle via structural changes in both thick and thin filaments. Proceedings of the National Academy of Sciences 113, E3039–E3047 (2016).

63. Lal, H. et al. Integrins and proximal signaling mechanisms in cardiovascular disease. Frontiers in bioscience : a journal and virtual library 14, 2307–2334 (2009).

64. Jacot, J.G. et al. Cardiac myocyte force development during differentiation and maturation. Ann N Y Acad Sci 1188, 121–127 (2010).

65. Lian, X. et al. Directed cardiomyocyte differentiation from human pluripotent stem cells by modulating Wnt/β-catenin signaling under fully defined conditions. Nature Protocols 8, 162–175 (2013).

66. Lian, X. et al. Robust cardiomyocyte differentiation from human pluripotent stem cells via temporal modulation of canonical Wnt signaling. Proceedings of the National Academy of Sciences 109, E1848–E1857 (2012).

67. Litviňuková, M. et al. Cells of the adult human heart. Nature 588, 466–472 (2020).

68. Avolio, E., Campagnolo, P., Katare, R. & Madeddu, P. The role of cardiac pericytes in health and disease: therapeutic targets for myocardial infarction. Nature Reviews Cardiology (2023).

69. Wang, R., Clark Ra Fau - Mosher, D.F., Mosher Df Fau - Ren, X.-D. & Ren, X.D. Fibronectin’s central cell-binding domain supports focal adhesion formation and Rho signal transduction. The Journal of biological chemistry (2005).

70. Zaidel-Bar, R., Milo, R., Kam, Z. & Geiger, B. A paxillin tyrosine phosphorylation switch regulates the assembly and form of cell-matrix adhesions. Journal of cell science 120, 137–148 (2007).

71. Deramaudt, T.B. et al. Altering FAK-Paxillin Interactions Reduces Adhesion, Migration and Invasion Processes. PLOS ONE 9, e92059 (2014).

72. Case, L.B. et al. Molecular mechanism of vinculin activation and nanoscale spatial organization in focal adhesions. Nature Cell Biology 17, 880–892 (2015).

73. Deakin, N.O., Ballestrem C Fau - Turner, C.E. & Turner, C.E. Paxillin and Hic-5 interaction with vinculin is differentially regulated by Rac1 and RhoA. PLOS ONE (2012).

74. Boopathy, G.T.K. & Hong, W. Role of Hippo Pathway-YAP/TAZ Signaling in Angiogenesis. Frontiers in cell and developmental biology (2019).

75. Werneburg, N., Gores, G.J. & Smoot, R.L. The Hippo Pathway and YAP Signaling: Emerging Concepts in Regulation, Signaling, and Experimental Targeting Strategies With Implications for Hepatobiliary Malignancies. Gene expression (2020).

76. Kwon, H., Kim, J. & Jho, E.A.-O. Role of the Hippo pathway and mechanisms for controlling cellular localization of YAP/TAZ. LID - 10.1111/febs.16091 [doi]. The FEBS journal (2021).

77. Teo, J., Lim, C.T., Yap, A. & Saw, T.B. A Biologist’s Guide to Traction Force Microscopy Using Polydimethylsiloxane Substrate for Two-Dimensional Cell Cultures. STAR Protocols 1, 100098 (2020).

78. Liu, J. et al. Talin determines the nanoscale architecture of focal adhesions. Proceedings of the National Academy of Sciences 112, E4864 (2015).

## SI References

1. C. J. Bult et al., Mouse Genome Database (MGD) 2019. Nucleic Acids Research 47, D801–D806 (2019).

